# Homocysteine Metabolites Impair the PHF8/H4K20me1/mTOR/Autophagy Pathway by Upregulating the Expression of Histone Demethylase PHF8-targeting microRNAs in Human Vascular Endothelial Cells and Mice

**DOI:** 10.1101/2023.06.27.546759

**Authors:** Łukasz Witucki, Hieronim Jakubowski

## Abstract

The inability to efficiently metabolize homocysteine (Hcy) due to nutritional and genetic deficiencies, leads to hyperhomocysteinemia (HHcy) and endothelial dysfunction, a hallmark of atherosclerosis which underpins cardiovascular disease (CVD). PHF8 is a histone demethylase that demethylates H4K20me1, which affects the mammalian target of rapamycin (mTOR) signaling and autophagy, processes that play important roles in CVD. PHF8 is regulated by microRNA (miR) such as miR-22-3p and miR-1229-3p. Biochemically, HHcy is characterized by elevated levels of Hcy, Hcy-thiolactone and *N*-Hcy-protein. Here, we examined effects of these metabolites on miR-22-3p, miR-1229-3p, and their target PHF8, as well as on the downstream consequences of these effects on H4K20me1, mTOR-, and autophagy-related proteins and mRNAs expression in human umbilical vein endothelial cells (HUVEC). We found that treatments with *N*-Hcy-protein, Hcy-thiolactone, or Hcy upregulated miR-22-3p and miR-1229-3p, attenuated PHF8 expression, upregulated H4K20me1, mTOR, and phospho-mTOR. Autophagy-related proteins (BECN1, ATG5, ATG7, lipidated LC3-II, and LC3-II/LC3-I ratio) were significantly downregulated by at least one of these metabolites. We also found similar changes in the expression of miR-22-3p, Phf8, mTOR- and autophagy-related proteins/mRNAs in vivo in hearts of *Cbs*^-/-^ mice, which show severe HHcy and endothelial dysfunction. Treatments with inhibitors of miR-22-3p or miR-1229-3p abrogated the effects of Hcy-thiolactone, *N*-Hcy-protein, and Hcy on miR expression and on PHF8, H4K20me1, mTOR-, and autophagy-related proteins/mRNAs in HUVEC. Taken together, these findings show that Hcy metabolites upregulate miR-22-3p and miR-1229-3p expression, which then dysregulate the PHF8/H4K20me1/mTOR/autophagy pathway, important for vascular homeostasis.

## 1 INTRODUCTION

Atherosclerosis, an inflammatory disease (1, 2), and its vascular manifestations, such as myocardial infarction, stroke, and peripheral artery disease, are the leading cause of morbidity and mortality worldwide. Endothelium supports vascular homeostasis by regulating the vascular tone and permeability. Endothelial dysfunction caused by biochemical or mechanical factors disrupts vascular homeostasis, induces inflammation, and is associated with atherothrombotic cardiovascular disease (CVD). The response-to-injury hypothesis proposes that endothelial dysfunction is the first step in atherosclerosis. In addition to elevated low-density lipoprotein, cigarette smoking, hypertension, diabetes mellitus, infectious microorganisms, and genetic alterations, endothelial dysfunction can be caused by elevated plasma homocysteine (Hcy), i.e., hyperhomocysteinemia (HHcy) (2). Indeed, endothelial dysfunction and inflammation are common findings in HHcy in humans and animal models, including Cbs-deficient mice (3-5). Inflammation is also seen in cultured endothelial cells treated with Hcy (6, 7) or Hcy-thiolactone (7).

Biochemically, HHcy is characterized by elevated levels of Hcy and its metabolites such as Hcy-thiolactone and *N*-Hcy-protein, seen in genetic and nutritional deficiencies in Hcy metabolism in humans and animals (8). Genetically or nutritionally induced HHcy causes pro-atherogenic changes in gene expression in mice and humans (9, 10).

Earlier studies have shown that human umbilical vein endothelial cells (HUVEC) metabolize Hcy to Hcy-thiolactone and *N*-Hcy-protein and that Hcy-thiolactone and *N*-Hcy-protein levels depend on extracellular concentrations of Hcy, folic acid, and HDL, factors that determine the susceptibility to vascular disease in humans, suggesting that Hcy-thiolactone and *N*-Hcy-protein could be involved in endothelial dysfunction and atherosclerosis (11). Later studies in HUVEC found that Hcy-thiolactone and *N*-Hcy-protein induce pro-atherogenic changes in gene expression that included upregulation by Hcy-thiolactone of LAMTOR2 mRNA, a part of the mammalian target of rapamycin (mTOR) signaling pathway (12). However, mechanisms by which these metabolites can affect gene expression in human vascular endothelial cells are not known.

MicroRNAs (miRs) are small non-coding RNAs which play a key role in cellular homeostasis by regulating gene expression at mRNA level (13). miR genes are transcribed by RNA polymerase II to primary miR, which is first cleaved by DROSHA and then by DICER (14, 15), generating 21 to 23-nucleotide long double strand mature miRs (16). One strand is incorporated into RISC (RNA-induced silencing complex) and regulates gene expression by binding to partially complementary mRNA sequence (mainly found in 3’ UTR) to induce translational repression, mRNA deadenylation or cleavage (17). Dysregulated miR expression can lead to endothelial dysfunction (18) and associated diseases such as CVD (19), and stroke (20).

Plant homeodomain finger protein 8 (PHF8) as one of the X chromosome genes linked to the intellectual disability syndrome, autism spectrum disorder, attention deficit hyperactivity disorder (21), and severe intellectual disability (22). PHF8 is a histone de-methylase that demethylates H4K20me1, which is important for homeostasis of the mTOR signaling (23). The expression of PHF8 is known to be regulated by microRNA (miR) such as miR-22-3p (24) and miR-1229-3p (25), which bind to PHF8 3’UTR. Studies in mice and neural cells showed that HHcy attenuated Phf8 expression (26). Whether HHcy affects PHF8 expression in human endothelial cells and whether miRs mediate effects of HHcy is unknown.

The mTOR signaling pathway is an important regulator of cellular metabolism and survival. When activated by nutrient abundance, it promotes anabolic processes such as protein biosynthesis and inhibits catabolic processes such as autophagy. Accumulating evidence suggests that mTOR signaling plays an important role in atherosclerosis and CVD (27, 28).

Autophagy is an evolutionarily conserved cellular process involving degradation and recycling damaged proteins and organelles that occurs continually at basal levels in cells and contributes to the maintenance of cellular homeostasis. Impaired autophagy flux causes the accumulation of damaged proteins and abnormal protein aggregates, and is associated with CVD, metabolic and neurodegenerative diseases (29).

We have previously demonstrated that HHcy activated mTOR signaling via a PHF8/H4K20me1-dependent mechanism and impaired autophagy in brains of mice genetically deficient in their ability to metabolize Hcy (*Cbs*^-/-^ mice (30, 31)) or Hcy-thiolactone (*Blmh*^-/-^ mice (32); *Pon1*^-/-^ mice (26)), and that Hcy metabolites such as Hcy-thiolactone and *N*-Hcy-protein activated mTOR signaling via the PHF8/H4K20me1-dependent mechanism, and impaired autophagy in cultured mouse neuroblastoma cells (26, 31, 32). However, mechanisms by which Hcy-thiolactone, *N*-Hcy-protein, or Hcy may affect mTOR- and autophagy-related proteins in human vascular endothelial cells were not known.

We hypothesize that HHcy-associated metabolites inhibit the expression of the histone demethylase PHF8 via a miR-mediated mechanism, which in turn influences mTOR- and autophagy-related proteins in human endothelial cells. To test this hypothesis, we studied how treatments with Hcy-thiolactone, *N*-Hcy-protein, or Hcy in HUVEC, and HHcy in Cbs-deficient mice, affect the expression of miRs targeting PHF8. We also examined how miR inhibitors influence the expression of PHF8 and its downstream targets.

## 2 MATERIALS AND METHODS

### 2.1 Cell culture and treatments

Human primary umbilical vein endothelial cells (HUVEC; ATCC, Manassas, VA, USA, Cat. # PCS-100-013) were grown in W2 flasks (37°C, 5% CO2) with 4mL of Vascular Cell Basal Medium (ATCC, Cat. # PCS-100-030) supplemented with 5% FBS and Endothelial Cell Growth Kit-VEGF (ATCC) and antibiotics (penicillin/streptomycin) (MilliporeSigma, Saint Louis, MO, USA). Cells between the 3rd and 10th passage were used in the experiments.

For experiments, 100,000 cells were seeded/each well of a 6-well plate. After cells reached 70-80% confluency (at 24 to 48 h), the monolayers were washed with PBS (2-times) and overlaid with M199 medium without methionine (Thermo Fisher Scientific, Waltham, MA, USA) having 5% dialyzed fetal bovine serum (FBS; Millipore Sigma). Cell cultures were treated with *N*-homocysteinylated-protein (*N*-Hcy-protein, prepared by modifying serum protein with Hcy-thiolactone as previously described, ref. (12)), L-Hcy-thiolactone, or D,L-Hcy (Millipore Sigma) and incubated at 37°C in 5% CO2 atmosphere for 24 h; cells from untreated cultures were used as controls. The concentrations of Hcy metabolites (*N*-Hcy protein, 10 and 20 µM; L-Hcy-thiolactone and D,L-Hcy, 20 and 200 µM) are based on their physiological concentrations (8) and earlier work (11) (12). These concentrations reflect levels of these metabolites in mice and humans (reviewed in ref. (8)).

For the miR inhibition experiments, HUVEC were transfected in Opti-MEM medium (Thermo Scientific) with miR inhibitors (Assay ID MH10203, for hsa-miR-22-3p or MH13382, for hsa-miR-1229-3p; Thermo Scientific) or mirVana™ miR Mimic, Negative Control #1 (Thermo Scientific, Cat. #4464058) as a control using Lipofectamine RNAiMax. Transformation efficiency was Target gene expression from the Negative Control-transfected cells is a baseline for evaluation of the effect of the experimental miRNA mimic on target gene expression. As suggested by the supplier, 0.5 nM miR or miR mimic, was used in the experiments. After 4-h-incubation, HUVEC were washed with PBS (2-times) and overlaid with M199 medium without methionine (Thermo Scientific) having 5% dialyzed FBS (MilliporeSigma). Cells were treated with D,L-Hcy, L-Hcy-thiolactone (Millipore Sigma), or *N*-Hcy-protein for 24 as above.

### 2.2 HUVEC viability assay

The viability of HUVEC was assessed by using the trypan blue exclusion assay based on the principle that the dye can enter membrane-compromised dead cells but is excluded by live cells. Briefly, 10,000 cells/well were seeded into wells of a 48-well plate. After cells reached 70-80% confluency (at 24-48 h), cells were treated as described in section 2.1 Cell culture and treatments. After 24 h, cells were rinsed with PBS (2-times), trypsinized, and pelleted by centrifuged (5 min, RT, 1,500 rpm). Cell pellets were suspended and incubated for 3 min in 350 µl PBS mixed with 150 µl 0.4% trypan blue solution (Millipore Sigma). For determining viability, cells were transferred to a hemocytometer chamber and counted using a light microscope at 10x magnification. The HUVEC viability is expressed as the percentage of blue-stained dead cells in the total number of cells.

### 2.3 Mice

Transgenic *Tg-I278T Cbs*^-/-^ mice on the C57BL/6J background harboring mutant human *CBS I278T* (*Tg-I278T*) transgene have been described before (33, 34). These mice are used as a model of human CBS deficiency, which is characterized by cardiovascular and neurological deficits (35). Briefly, in these mice, the expression of human mutant CBS, which has less than 3% of the wild-type CBS enzyme activity, is controlled by the zinc-inducible metallothionein promoter, which allows to rescue the neonatal lethality phenotype of *Cbs*^-/-^ in mice by adding 25 mM ZnCl2 to the drinking water of pregnant dams. Zinc-water is replaced by plain water after weaning at 30 days. *Cbs*^-/-^ mice, but not *Cbs*^+/-^ heterozygotes, exhibit reduced body weight, increased susceptibility to atherosclerosis, and shortened life span compared to *Cbs*^+/-^ mice which are indistinguishable from *Cbs*^+/+^ animals. *Cbs*^-/-^ mice have severely elevated total Hcy (plasma 272±50 μM, urine 4.1±1.4 mM), *N*-Hcy-protein (plasma 16.6±4.1μM; urine10.8±4.1 μM), and Hcy-thiolactone (urine 11.8±0.9 μM) compared to *Cbs*^+/-^ (plasma tHcy 5.0±2.6 μM and *N*-Hcy-protein, 2.6±1.7 μM; urine Hcy-thiolactone, <0.2 nM) and *Cbs*^+/+^ animals (plasma tHcy 3.4±0.3 μM and *N*-Hcy-protein 1.9±0.7 μM; Hcy-thiolactone, urine <0.2 nM).

Mouse *Cbs* genotypes were established by PCR using the following primers: *forward 5′-*GGTCTGGAATTCACTATGTAGC*-3′*, wild type reverse *5′-*CGGATGACCTGCATTCATCT*-3′,* mutant reverse: *5′-*GAGGTCGACGGTATCGATA*-3′.* The mice, fed with a standard rodent chow (LabDiet5010; Purina Mills International, St. Louis MO, USA), were maintained at the Rutgers-New Jersey Medical School Animal Facility. Twelve-month-old *Cbs*^-/-^ female mice and their *Cbs*^+/-^ female siblings as controls were used in experiments. Animal procedures were approved by the Institutional Animal Care and Use Committee at Rutgers-New Jersey Medical School.

### 2.4 Protein extraction

#### 2.4.1 HUVEC

Proteins were extracted from 300,000 to 400,00 HUVEC cells (per one well of a 6-well plate) with RIPA buffer (Millipore Sigma) according to manufacturer’s protocol, quantified using PierceTM BCA protein Kit (Termo Fisher).

#### 2.4.2 Mouse hearts

Mice were killed by CO2 inhalation, the hearts collected, frozen on dry ice, pulverized with dry ice using a mortar and pestle, and stored at −80°C. Proteins were extracted from the pulverized hearts (50±5 mg) using RIPA buffer (4 v/w, containing protease and phosphatase inhibitors) with sonication (Bandelin SONOPLUS HD 2070) on wet ice (three sets of five 10-s strokes with 1 min cooling interval between strokes). Brain extracts were clarified by centrifugation (15,000 g, 15 min, 4°C) and clear protein extracts containing 8-12 mg protein/mL were collected. Protein concentrations were quantified using the BCA kit (Thermo Scientific).

### 2.5 Western blots

Proteins were separated by SDS-PAGE on 10% gels (8 µg protein/lane, 5 % total), and transferred to PVDF membrane (Bio-Rad) for 20 min at 0.1 A, 25 V using Trans Blot Turbo Transfer System (Bio-Rad) as previously described (26, 36). After blocking with 5 % bovine serum albumin in TBST buffer (1h, room temperature), the membranes were incubated with anti-PHF8 (Abcam, ab36068), anti-H4K20me1 (Abcam ab177188), anti-mTOR (CS #2983), anti-pmTOR Ser2448 (CS, #5536), anti-ATG5 (CS, #12994), anti-ATG7 (CS, #8558), anti-BECN1 (CS, #3495), anti-p62 (CS, #39749), anti-LC3 (CS, #12741), anti-GAPDH (CS, #5174) for 16 hours. Membranes were washed three times with 1X Tris-Buffered Saline, 0.1% Tween 20 Detergent (TBS-T), 10 min each, and incubated with goat anti-rabbit IgG secondary antibody conjugated with horseradish peroxidase. Positive signals were detected using Western Bright Quantum-Advansta K12042-D20 and GeneGnome XRQ NPC chemiluminescence detection system. Bands intensity was calculated using Gene Tools program from Syngene. GAPDH protein was used as a reference for quantification.

### 2.6 mRNA and miR quantification by RT-qPCR

Total RNA was isolated using Trizol reagent (Millipore Sigma). cDNA synthesis was conducted using Revert Aid First cDNA Synthesis Kit (Thermo Fisher Scientific) according to manufacturer’s protocol. Nucleic acid concentration was measured using NanoDrop (Thermo Fisher Scientific). RT-qPCR was performed with SYBR Green Mix and CFX96 thermocycler (Bio-Rad). GAPDH mRNA was used as a reference for mRNA quantification. RT-qPCR primer sequences are listed in Supplementary Table S1.

One μg of total RNA was used for the microRNA polyadenylation and re-verse-transcription reactions. MicroRNAs (miRs) were polyadenylated and then re-verse-transcribed using the miR 1st-Strand cDNA Synthesis Kit (Agilent Technologies) according to manufacturer’s protocol.

cDNA was used for RT-qPCR with miRNA QPCR Master Mix (Agilent Technologies). Universal reverse primers (Agilent Technologies) and unique miR-specific primers (same sequence as an analyzed miR) (Table 1) were used to quantify miR levels. Reactions were conducted on CFX96 thermocycler (Bio-Rad). 18S rRNA and U6 snRNA were used as references for miR quantification.

The 2(-ΔΔCt) method was used to calculate the relative expression levels (37). Data analysis was performed with the CFX Manager™ Software, Microsoft Excel, and GraphPad Prism7.

### 2.7 Dual luciferase assay

The 3′UTR fragment of the human *PHF8* gene having native or mutated miR-22-3p or miR-1229-3p binding sites (Supplementary Figure S1) was cloned into the pmirGLO vector (Promega) cut with XbaI and DraI restriction enzymes (New England BioLabs). Transformation efficiency was 50% for plasmids containing a mir-22 target site and 75% for mir-1229 target site. To confirm miR-PHF8 interaction, we performed dual luciferase reporter assays according to manufacturer’s protocol. Briefly, 15,000 HUVEC cells were seeded on each well on 96 well plates, grown to 70% confluency (^˷^24 h), and transfected with a *3’UTR PHF8*-containg plasmid (0.01 µg) in the absence and presence of a specific miR inhibitor (0.5 nM) using Lipofectamine 2000 (Invitrogen). After 4-h, HUVEC monolayers were washed twice with PBS, overlaid with M199 medium without methionine (Thermo Scientific) containing 5% dialyzed FBS (Millipore Sigma), and treated with D,L-Hcy, L-Hcy-thiolactone (Millipore Sigma) or *N*-Hcy-protein for 24 h. The firefly and renilla luminescence were quantified using a Dual-Glo® Luciferase Assay System (Promega, USA) and the firefly/renilla luminescence ratios calculated.

### 2.8 Statistical analyses

Each assay was repeated three times (technical repeats) in three independent experiments (biological repeats) for each treatment and controls. Data as mean ± standard deviation (SD) of three biological repeats. Values for each experimental/treatment group were normalized to controls. Data were analyzed using the Kruskal-Wallis test was used for comparisons of more than two groups and the Mann-Whitney or *T*-test for comparisons of two groups. The analyses were carried out using GraphPad Prism7 software (GraphPad Holdings LLC, San Diego CA, USA, https://www.graphpad.com).

## 3 RESULTS

### 3.1 Hcy-thiolactone, *N*-Hcy-protein, and Hcy downregulate the histone demethylase PHF8, upregulate the H4K20me1 epigenetic mark and mTOR, and impair autophagy in HUVEC

HUVEC can metabolize Hcy to Hcy-thiolactone and *N*-Hcy-protein (11). To figure out how each of these metabolites affects the expression of PHF8 and its effects on downstream targets, we treated HUVECs with Hcy-thiolactone, *N*-Hcy-protein, and Hcy. We found significantly reduced expression of PHF8 protein, quantified by western blotting, in HUVECs treated with Hcy-thiolactone or *N*-Hcy-protein while Hcy tended to reduce PHF8 levels compared to control (Figure 1A).

**Figure 1.**
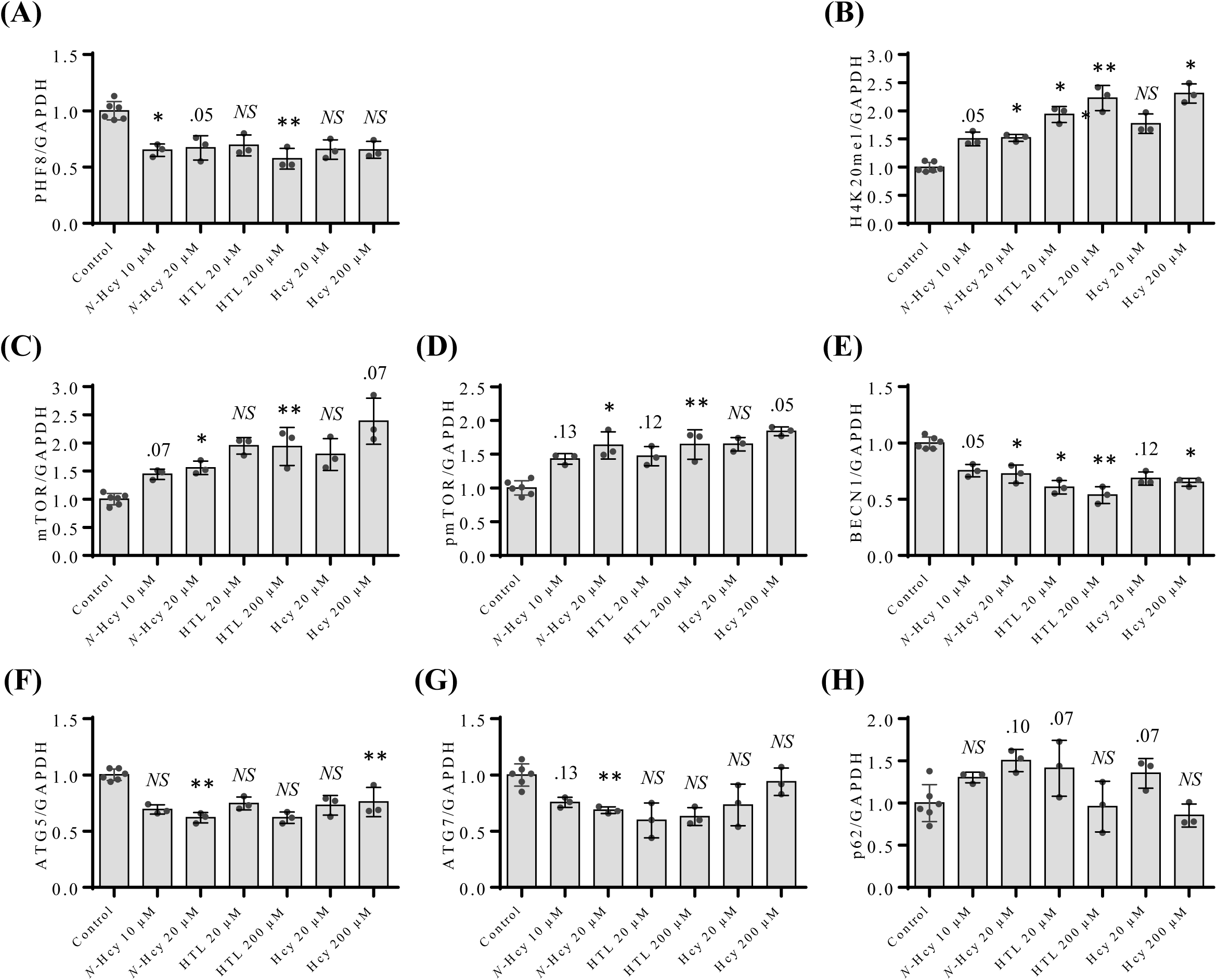
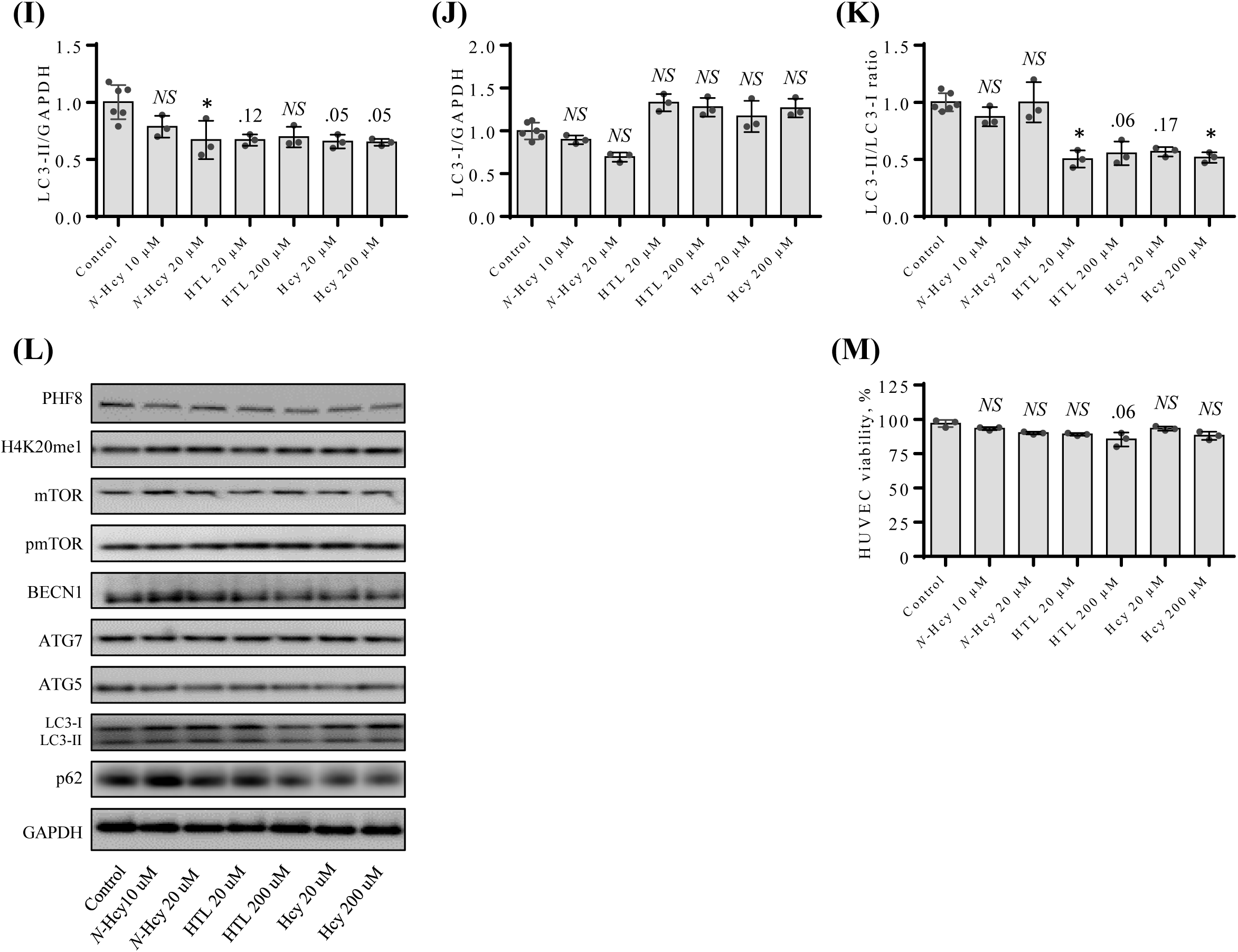
*N*-Hcy-protein, Hcy-thiolactone, and Hcy downregulate PHF8, upregulate the H4K20me1 epigenetic mark, mTOR signaling, and impair autophagy in HUVEC. The cells were treated with 20 or 200 µM of *N*-Hcy-protein (N-Hcy) or Hcy-thiolactone (HTL) for 24 h at 37⁰C. Untreated cells were used as controls. Bar graphs illustrating the quantification of PHF8 (A), H4K20me1 (B), mTOR (C), pmTOR (D), BECN1 (E), ATG5 (F), ATG7 (G), p62 (H), LC3-I (I), LC3-II (J), LC3-I/LC3-II ratio (K) based on western blot analyses are shown. GAPDH was used as a reference protein. Representative images of western blots are shown in panel (L). The treatments with HHcy-related metabolites did not affect HUVEC viability (M). Each assay was repeated three times (technical repeats) in three independent experiments (biological repeats). Mean SD values for each treatment group are shown. *P*-values were calculated by Kruskal-Wallis nonparametric test. * *P* < 0.05, ** *P* < 0.01. The numbers above bars show P values 0.05 – 0.17. NS, not significant; *N*-Hcy, *N*-Hcy-protein.

We also found significantly elevated levels of the methylated histone H4K20me1, a positive epigenetic regulator of mTOR expression, in HUVEC treated with *N*-Hcy-protein, Hcy-thiolactone, or Hcy, compared to untreated control cells (Figure 1B).

The expression of mTOR protein was significantly upregulated in HUVEC by treatments with these metabolites (Figure 1C). As mTOR is activated by phosphorylation, we quantified the active form of mTOR, phosphorylated at Ser2448 (pmTOR) and found that pmTOR was also significantly upregulated by treatments with N-Hcy-protein or Hcy-thiolactone while Hcy tended to upregulate pmTOR compared to control (Figure 1D).

The regulators of autophagosome assembly BECN1, ATG5, and ATG7 proteins were significantly downregulated (Figure 1E, 1F, and 1G, respectively) in HUVEC treated with *N*-Hcy-protein. Treatments with Hcy-thiolactone or Hcy downregulated BECN1 (Figure 1E), while Hcy downregulated ATG5 (Figure 1F) with the effects becoming significant at higher concentrations.

To find out whether HHcy-related metabolites affect the autophagy flux, we also quantified microtubule-associated protein 1 light chain 3 (LC3) and p62 protein, a receptor for degradation of ubiquitinated substrates. We found reductions in lipidated LC3-II (Figure 1I) in HUVEC treated with *N*-Hcy-protein, Hcy-thiolactone, or Hcy while unlipidated LC3-I was not significantly affected (Figure 1J). The LC3-II/LC3-I ratio, a measure of the autophagy flux, was significantly reduced in cells treated with Hcy-thiolactone or Hcy but not *N*-Hcy-protein (Figure 1K), while p62 protein was not significantly affected in cells treated with these metabolites (Figure 1H). These findings suggest that Hcy metabolites can impair autophagy flux in a metabolite-specific manner. Representative images of western blots are shown in Figure 1L.

Cell viability was not significantly affected in HUVEC treated with *N*-Hcy-protein, Hcy-thiolactone, or Hcy, compared to untreated control (Figure 1M).

These findings show that Hcy and its downstream metabolites, Hcy-thiolactone and *N*-Hcy-protein, can influence the expression of PHF8 and its downstream targets, mTOR signaling-related and autophagy-related proteins.

### 3.2 Hcy-thiolactone, *N*-Hcy-protein, and Hcy downregulate PHF8 by upregulating the expression of miR-22-3p and miR-1229-3p targeting PHF8

To elucidate the mechanism by which Hcy metabolites affect PHF8 expression we first quantified PHF8 mRNA by RT-qPCR to find out whether Hcy metabolites exert transcriptional control on PHF8 expression in HUVECs. We found that PHF8 mRNA was significantly downregulated in HUVECs treated with *N*-Hcy-protein, Hcy-thiolactone, or Hcy, compared to untreated control (Figure 2A), reflecting similar downregulation seen in the PHF8 protein levels (Figure 1A). These findings show that the effects of Hcy metabolites on PHF8 expression are transcriptional. Methionine did not affect PHF8 mRNA (not shown).

**Figure 2.**
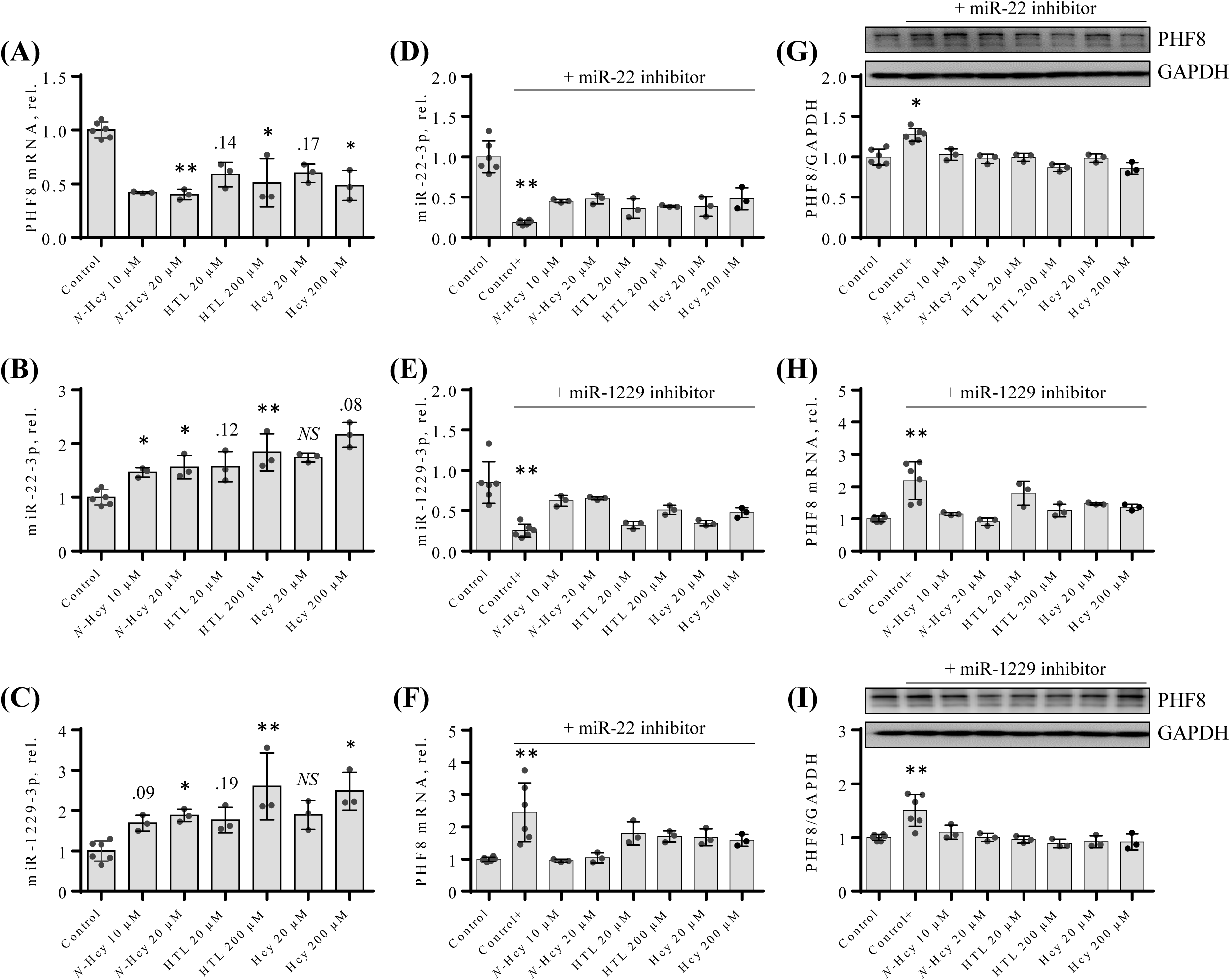
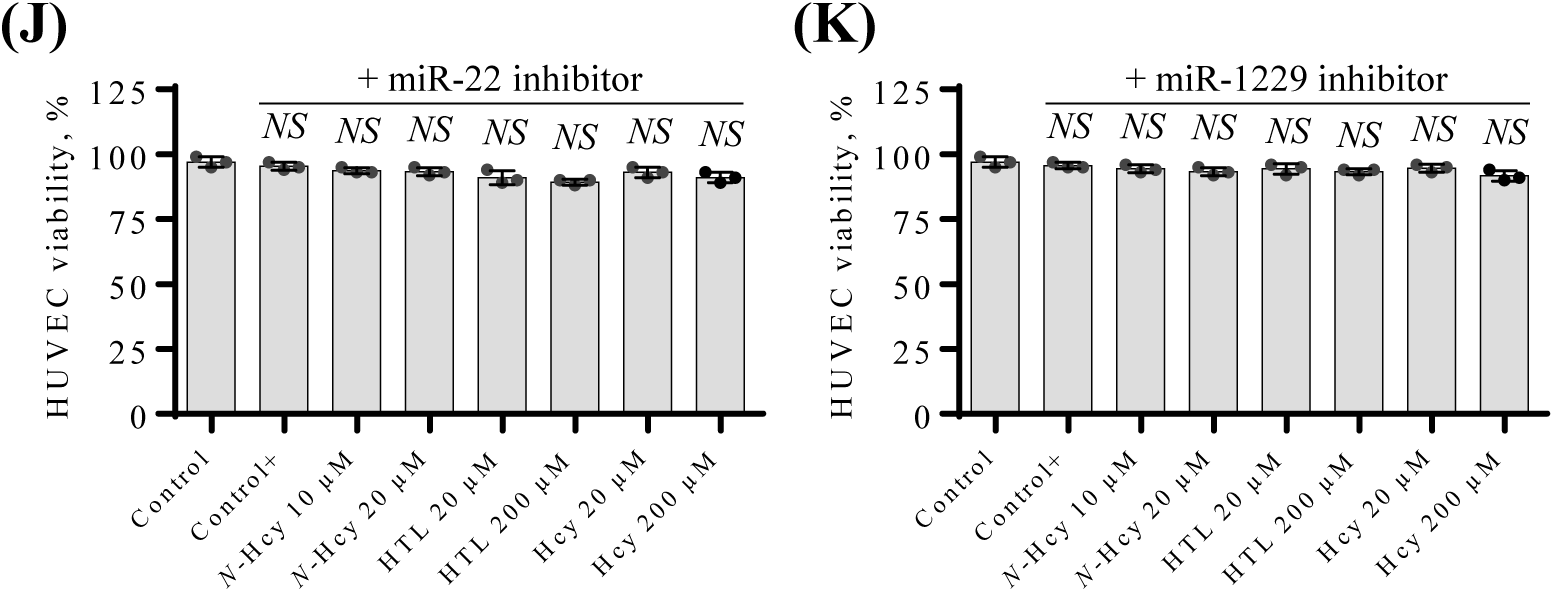
Effects of *N*-Hcy-protein, Hcy-thiolactone, and Hcy on the expression of PHF8 mRNA, miR-22-3p, and miR-1229-3p in the absence (A – C) or presence (D - I) of miR inhibitors. (A - C) HUVEC were treated with *N*-Hcy-protein (N-Hcy), Hcy-thiolactone (HTL), or Hcy for 24 h and PHF8 mRNA (A), miR-22-3p (B), and miR-1229-3p (C) were quantified by RT-qPCR. Untreated cells were used as controls. (D - I) HUVEC were transfected with were transfected with Thermo Scientific mirVana™ miRNA Mimic, Negative Control #1 (Control), inhibitor of miR-22-3p (D, F, G), or inhibitor of miR-1229-3p (E, H, I) for 4 h. The cells transfected with a miR inhibitor were then untreated (Control+) or treated with N-Hcy-protein, Hcy-thiolactone, or Hcy in methionine-free M199/dialyzed FBS medium for 24 h. The expression of miR-22-3p (D), and miR-1229-3p (E), and PHF8 mRNA (F, H) was quantified by RT-qPCR. GAPDH mRNA was used as a reference for PHF8 mRNA. 18S rRNA and U6 snRNA were used as references for miR quantification. Bar plots in panels (G) and (I) show the expression of PHF8 protein quantified by western blotting. GAPDH was used as a reference protein. Representative images of western blots are shown above the bar plots in panels (G) and (I). The transfections with miR inhibitors and treatments with HHcy-related metabolites did not affect HUVEC viability (L, M). Each assay was repeated three times (technical repeats) in three independent experiments (biological repeats). Mean SD values for each treatment group are shown. P-values were calculated by Kruskal-Wallis nonparametric test (A – C) or Mann Whitney test (D - I). * *P* < 0.05, ** *P* < 0.01. The numbers above bars show *P*-values from 0.06 to 0.19. NS, not significant; *N*-Hcy, *N*-Hcy-protein.

To find out whether the transcriptional downregulation of PHF8 caused by Hcy metabolites is mediated by PHF8 mRNA-targeting miRs in HUVECs, we quantified by RT-qPCR miR-22-3p and miR-1229-3p that have been previously suggested to bind to the PHF8 3’UTR. Target site for miR-22-3p on the PHF8 3’UTR hs been confirmed by the dual-luciferase reporter vector assay in gastric cancer cell lines (ref. 24; Supplementary Figure S1). Target site for miR-1229-3p on the PHF8 3’UTR has been suggested by analyses of miR-mRNA specific interaction using the crosslinking, ligation, and sequencing of hybrids in human embryonic kidney Flp-In T-REx 293 cells (25).

We found significantly upregulated miR-22-3p levels in HUVECs treated with *N*-Hcy-protein or Hcy-thiolactone compared to untreated control (Figure 2B). Hcy did not significantly affect miR-22-3p levels. Methionine did not affect miR-22-3p expression (not shown).

We also found significantly upregulated miR-1229-3p levels in HUVEC treated with N-Hcy-protein, Hcy-thiolactone, or Hcy compared to control (Figure 2C). Methionine did not affect miR-1229-3p expression (not shown).

### 3.3 Treatments with miR inhibitors ameliorate effects of Hcy-thiolactone, *N*-Hcy-protein, and Hcy on miR-22-3p, miR-1229-3p, and PHF8 expression

To verify these findings, we carried out experiments with miR inhibitors. We found that transfections of HUVECs with miR-22-3p inhibitor significantly reduced miR-22-3p levels (to 17% compared to untreated control; Figure 2D) while transfections with miR-1229-3p inhibitor reduced miR-1229-3p levels (to 22% compared to control; Figure 2E).

We also found that treatments of HUVEC with miR-22-3p inhibitor significantly increased PHF8 mRNA (Figure 2F) and PHF8 protein level (Figure 2G). Treatments with miR-1229 inhibitor significantly increased PHF8 mRNA (Figure 2H) and protein level (Figure 2I).

At the same time, inhibitors of miR-22-3p and miR-1229-3p ameliorated stimulatory effects of Hcy-thiolactone, *N*-Hcy-protein, and Hcy on miR-22-3p (Figure 2D) and miR-1229-3p expression (Figure 2E) seen in the absence of these inhibitors (Figure 2B and Figure 2C, respectively).

Inhibitors of miR-22-3p and miR-1229 also ameliorated inhibitory effects of Hcy-thiolactone, *N*-Hcy-protein, and Hcy on PHF8 mRNA (Figure 2F and Figure 2H, respectively) and PHF8 protein expression (Figure 2G and Figure 2I, respectively) seen in the absence of these inhibitors (PHF8 mRNA, Figure 2A; PHF8 protein, Figure 1A). Representative images of western blots are shown above the bar plots in Figure 2G, I.

Cell viability was not significantly affected in HUVEC transfected with miR-22-3p inhibitor (Figure 2J) or miR-1229-3p inhibitor (Figure 2K) and treated with *N*-Hcy-protein, Hcy-thiolactone, or Hcy, compared to untreated control.

### 3.4 Dual luciferase assays

In dual luciferase reporter assays, the activity of PHF8 3’UTR containing native binding sites for miR-1229-3p or miR-22-3p was significantly inhibited by Hcy-thiolactone, Hcy, or *N*-Hcy-protein (panels A and C, Supplementary Figure S2) consistent with the effects of these metabolites on the expression of PHF8 protein (Figure 1A) and mRNA (Figure 2A) in HUVEC. The inhibitory effects of Hcy-thiolactone, Hcy, or *N*-Hcy-protein were also not seen in experiments with PHF8 3’UTR containing mutated binding sites for miR-1229-3p or miR-22-3p (panels B and D, respectively, Supplementary Figure S2). These inhibitory effects of Hcy-thiolactone, Hcy, or *N*-Hcy-protein on PHF8 3’UTR-dependent luciferase activity were abrogated by inhibitors of miR-1229-3p or miR-22-3p (panels C and E, respectively, Supplementary Figure S2), thus recapitulating effects of these miR inhibitors on the expression of PHF8 protein (Figure 2G) and mRNA (Figure 2H) in HUVEC. The inhibitors of miR-1229-3p or miR-22-3p stimulated the activity of native but not mutant PHF8 3’UTR also in the absence of Hcy metabolites (Supplementary Figure 3).

Taken together, these findings confirm that PHF8 3’UTR is a target for miR-1229-3p and miR-22-3p.

### 3.5 Effects of miR inhibitors are propagated to downstream targets of PHF8

As PHF8 is a master regulator of mTOR expression, which in turn influences autophagy, we predicted that the primary effect of miR-22-3p and miR-1229-3p inhibitors on PHF8 will be propagated to targets downstream of PHF8. To examine this prediction, we quantified mTOR- and autophagy-related mRNAs and proteins in HUVEC transfected with miR inhibitors. We found that the miR-22-3p inhibitor significantly downregulated the expression of mTOR mRNA (Figure 3A). Autophagy-related ATG5, ATG7, and BECN1 mRNAs were significantly upregulated (Figure 3B, Figure 3C, and Figure 3D, respectively). Levels of LC3 mRNA and p62 mRNA were not significantly affected by the miR-22-3p inhibitor (not shown).

**Figure 3.**
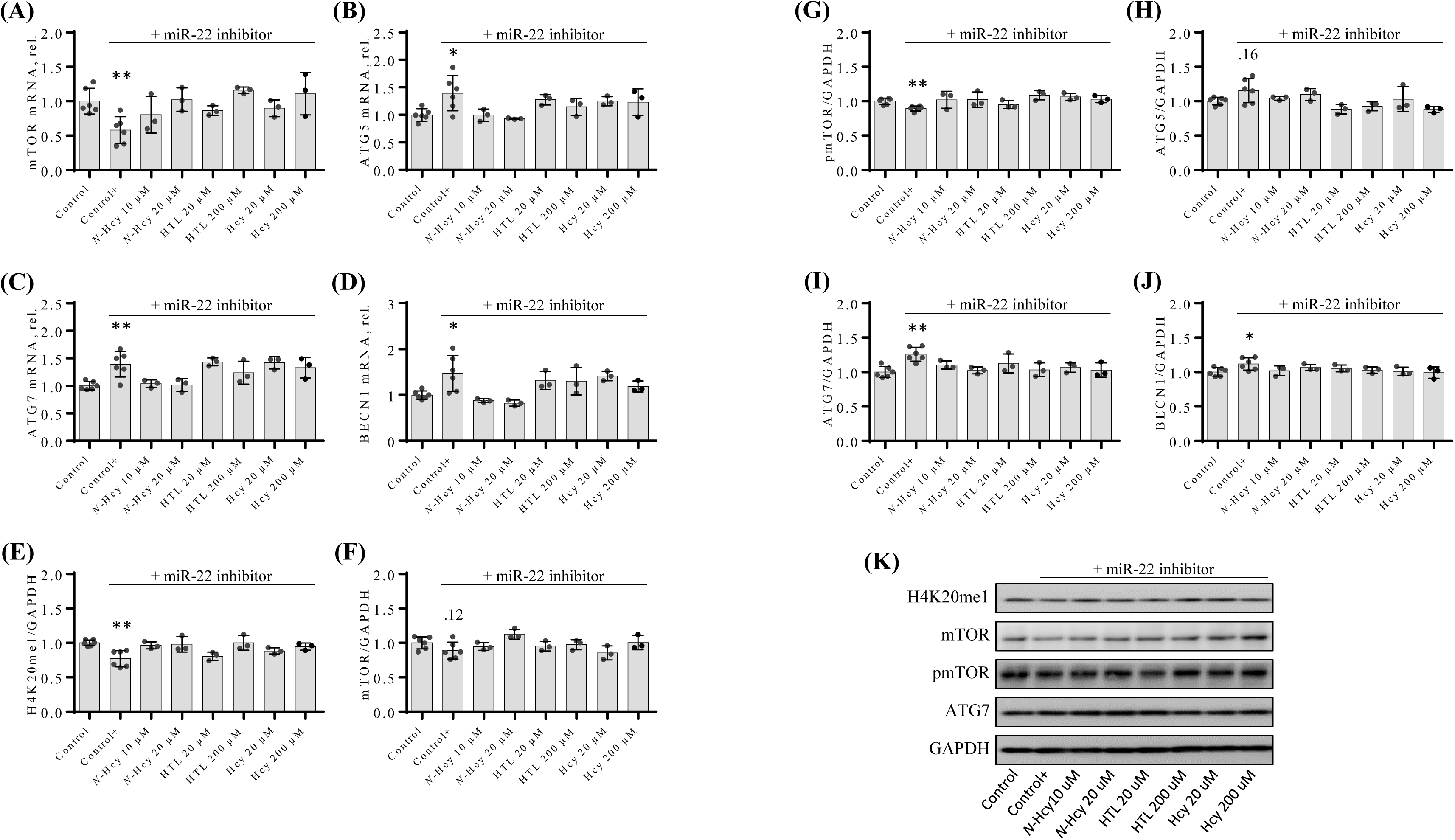
The inhibitor of PHF8-targeting miR-22-3p ameliorates the influence of Hcy metabolites on the expression of mTOR- and autophagy related mRNAs (A - D) and proteins (E - K). HUVEC were transfected with Thermo Scientific mirVana™ miRNA Mimic, Negative Control #1 (Control) or the inhibitor of miR-22-3p for 4 h. The cells transfected with the miR inhibitor were then un-treated (Control+) or treated with *N*-Hcy-protein (N-Hcy), Hcy-thiolactone (HTL), or Hcy in methionine-free M199/dialyzed FBS medium for 24 h. The indicated mRNAs and proteins were quantified by RT-qPCR and western blotting, respectively. GAPDH mRNA was used as a reference for mTOR (A), ATG5 (B), ATG7 (C), and BECN1 (D) mRNAs. GAPDH protein was used as a reference for H4K20me1 (E), mTOR (F), pmTOR (G), ATG5 (H), ATG7 (I), and BECN1 (J) proteins. Representative images of western blots are shown in panel (K). Each assay was repeated three times (technical repeats) in three independent experiments (biological repeats). Mean SD values for each treatment group are shown. *P*-values were calculated by the Mann-Whitney test. * *P* < 0.05, ** *P* < 0.01. The numbers above bar in (F) and (H) are *P*-values >0.05. NS, not significant; *N*-Hcy, *N*-Hcy-protein.

The effect of miR-22-3p inhibitor on PHF8 was also propagated to mTOR- and autophagy-related proteins. Specifically, the miR-22-3p inhibitor significantly downregulated H4K20me1 (Figure 3E), mTOR (Figure 3F), and pmTOR (Figure 3G) protein levels compared to control. Autophagy-related proteins ATG7 (Figure 3I) and BECN1 (Figure 3J) were significantly upregulated while ATG5 protein was not affected by miR-22-3p inhibitor (Figure 3H). Representative images of western blots are shown in Figure 3K.

The effects of miR-1229 inhibitor on PHF8 were also propagated to mTOR- and autophagy-related mRNAs and proteins. Specifically, miR-1229 inhibitor significantly downregulated the expression of mTOR mRNA (Figure 4A) and significantly upregulated autophagy-related mRNAs such as ATG5 (Figure 4B), ATG7 (Figure 4C), and BECN1 (Figure 4D). LC3 mRNA appeared not to be affected (Figure 4E) and p62 mRNA was not affected (Figure 4F) by the miR-1229 inhibitor.

**Figure 4.**
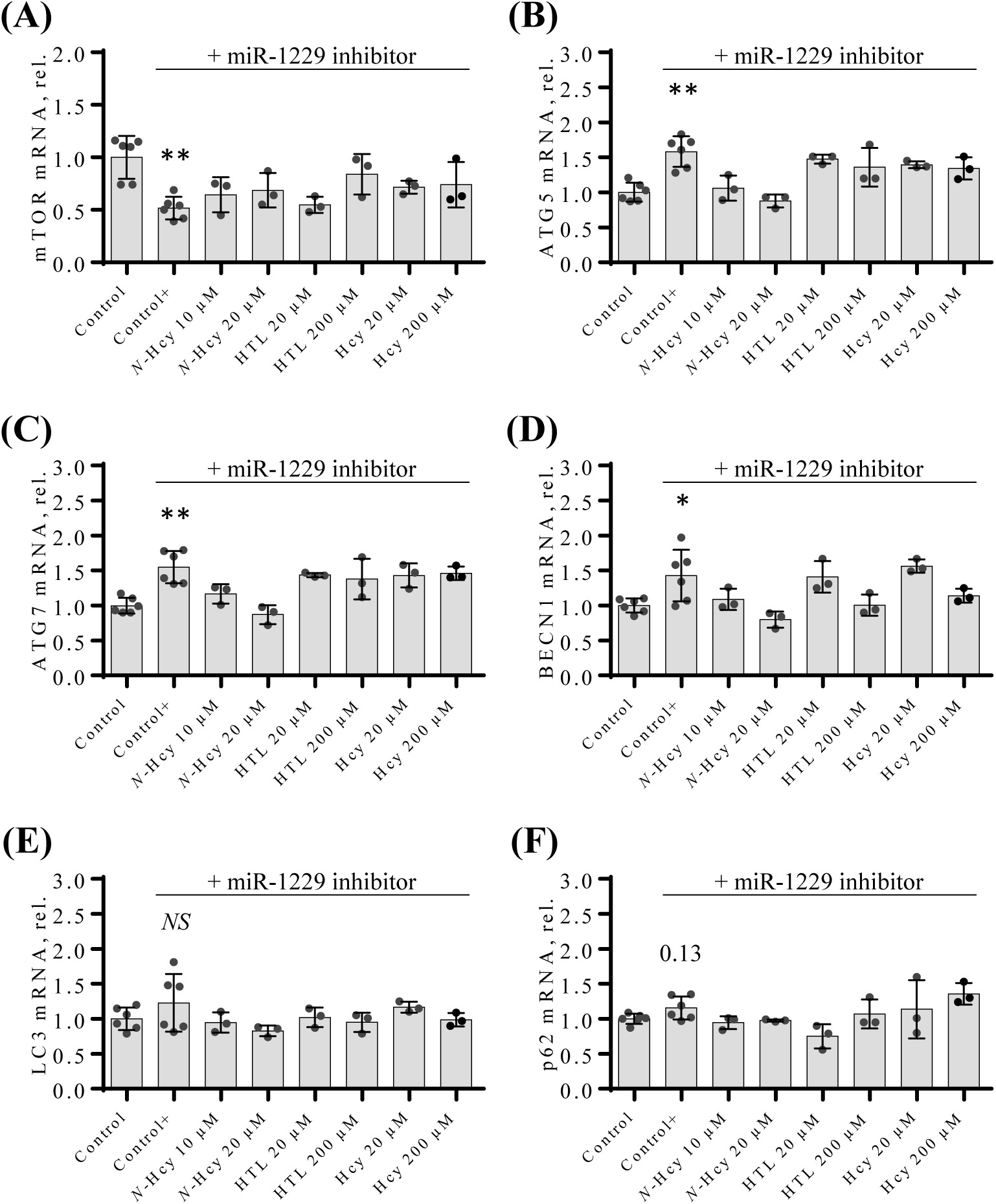
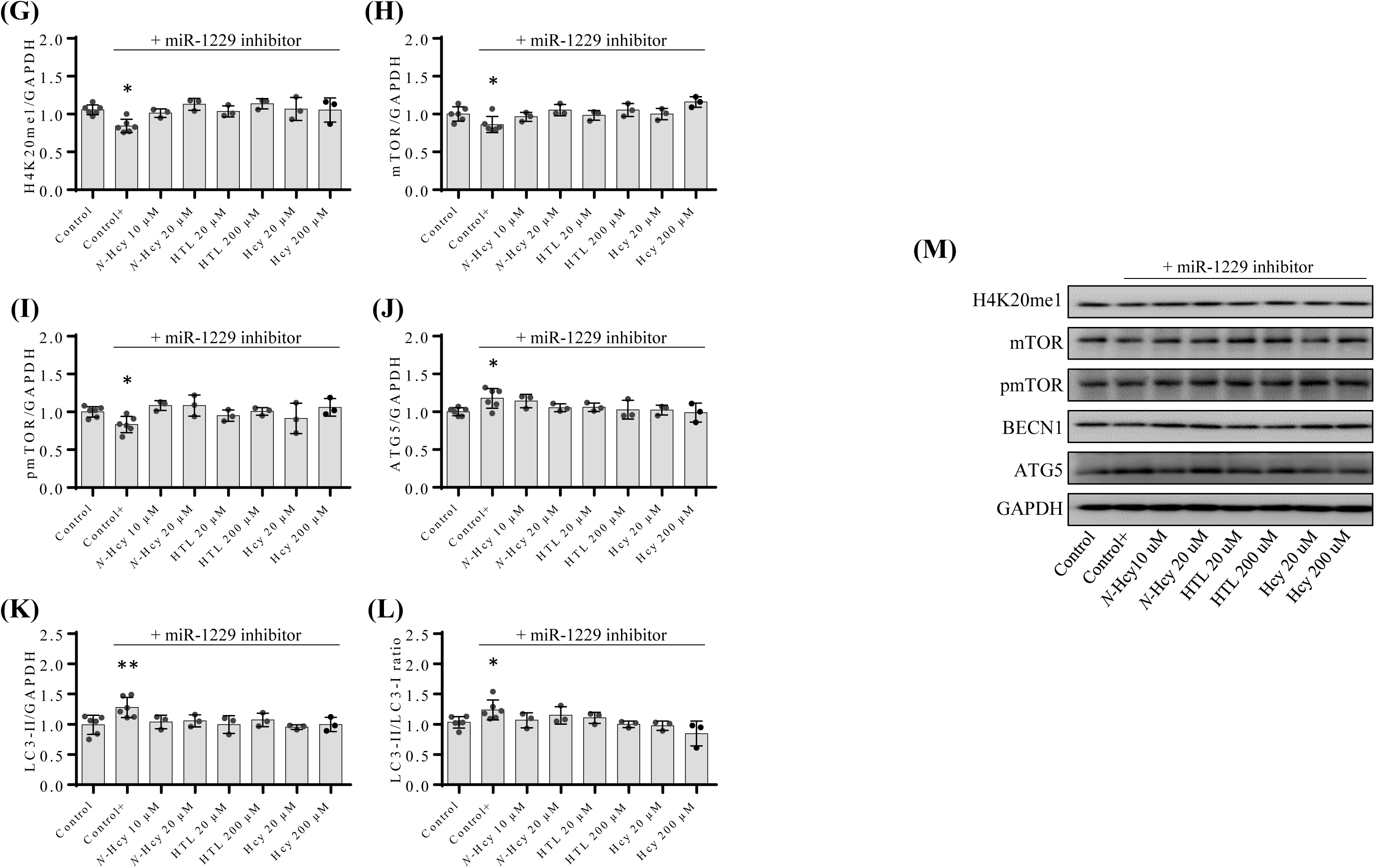
The inhibitor of PHF8-targeting miR-1229-3p ameliorates the influence of Hcy metabolites on the expression of mTOR- and autophagy related mRNAs (A - F) and proteins (G - L). HUVEC were transfected with Thermo Scientific mirVana™ miRNA Mimic, Negative Control #1 (Control) or the inhibitor of miR-1229-3p for 4 h. The cells transfected with the miR inhibitor were then untreated (Control+) or treated with *N*-Hcy-protein (*N*-Hcy), Hcy-thiolactone (HTL), or Hcy for 24 h. The indicated mRNAs were quantified by RT-qPCR and proteins by western blotting. GAPDH mRNA was used as a reference for mTOR (A), ATG5 (B), ATG7 (C), BECN1 (D), LC3 (E), and p62 (F) mRNAs. GAPDH protein was used as a reference for H4K20me1 (G), mTOR (H), pmTOR (I), ATG5 (J), and LC3-II (K) proteins. Panel (L) shows the LC3-II/LC3-I ratio. Representative images of western blots are shown in panel (M). Each assay was repeated three times (technical repeats) in three independent experiments (biological repeats). Mean SD values for each treatment group are shown. P-values were calculated by the Mann Whitney test. * *P* < 0.05, ** *P* < 0.01. The number 0.13 above the ‘Control+’ bar in (F), is a *P*-value. NS, not significant; *N*-Hcy, *N*-Hcy-protein.

Proteins such as H4K20me1 (Figure 4G), mTOR (Figure 4H), and pmTOR (Figure 4I) were significantly downregulated by the miR-1229 inhibitor. Autophagy-related proteins ATG5 (Figure 4J), lipidated LC3-II (Figure 4K), and LC3-II/LC3-I ratio (P < 0.05, Figure 4L) were significantly upregulated, while LC3-I was not affected (not shown), by miR-1229 inhibitor. Representative images of western blots are shown in Figure 4M.

### 3.6. Treatments with miR inhibitors ameliorate effects of Hcy-thiolactone, *N*-Hcy-protein, and Hcy on mTOR- and autophagy-related mRNA and protein expression

We also found that treatments of HUVEC with inhibitors of miR-22-3p and miR-1229-3p ameliorated stimulatory effects of Hcy-thiolactone, *N*-Hcy-protein, and Hcy on H4K20me1 (Figure 3E and 4G), mTOR mRNA (Figure 3A and 4A), mTOR protein (Figure 3F and 4H), and pmTOR protein (Figure 3G and 4I) seen in the absence of these inhibitors (Figure 1B, 1C, 1D).

Inhibitor of miR-22-3p ameliorated inhibitory effects of Hcy-thiolactone, N-Hcy-protein, and Hcy on ATG5, ATG7, and BECN1 mRNA (Figure 3B, 3C, and 3D, respectively) and protein (Figure 3H, 3I, and 3J, respectively) seen in the absence of these inhibitors (Figure 1E, 1F and 1G, respectively). Direct comparisons of Figure 3 (+inhibitor) and Figure 1 (-inhibitor) data are shown in Supplementary Table S2.

Inhibitor of miR-1229 ameliorated inhibitory effects of Hcy-thiolactone, *N*-Hcy-protein, and Hcy on ATG5, ATG7, BECN1, and LC3 mRNA (Figure 4B, 4C, 4D, and 4E, respectively), ATG5 protein (Figure 4J), LC3-II protein (Figure 4K), and LC3-II/LC3-1 ratio (Figure 4L). Representative images of western blots are shown in Figure 3K and Figure 4M.

### 3.7 Cbs deficiency upregulates miR-22-3p and dysregulates the Phf8/H4K20me3/mTOR/ autophagy pathway in the mouse heart

Endothelial dysfunction is commonly seen in murine models of HHcy, including Cbs-deficient mice (3-5), but whether miR-22-3p is involved was not known. We studied miR-22-3p expression in *Cbs*^-/-^ mice that are characterized by severely elevated levels of Hcy, Hcy-thiolactone, and *N*-Hcy-protein, metabolites which we found to upregulate miR-22-3p (Figure 2B) and miR-1229-3p (Figure 2C) and downregulate PHF8 protein (Figure 1A) and (Figure 2B) in HUVEC.

To find out if miR-dependent regulation of Phf8 expression occurs in vivo, we quantified miR-22-3p (miR-1229 is not found in mice) and Phf8 in hearts of 12-month-old *Tg-I278T Cbs*^-/-^ mice and *Tg*-*I278T Cbs*^+/-^ sibling controls. We found that miR-22-3p was significantly upregulated in hearts of *Tg*-*I278T Cbs*^-/-^ mice compared to *Tg*-*I278T Cbs*^+/-^ sibling controls (*P* < 0.05, Figure 5A). At the same time, levels of Phf8 mRNA (*P* < 0.01; Figure 5A) and protein (*P* < 0.01; Figure 5B) were significantly downregulated. These findings show that miR-22-3p is a negative regulator of Phf8 expression in hearts of *Cbs*^-/-^ mice.

**Figure 5.**
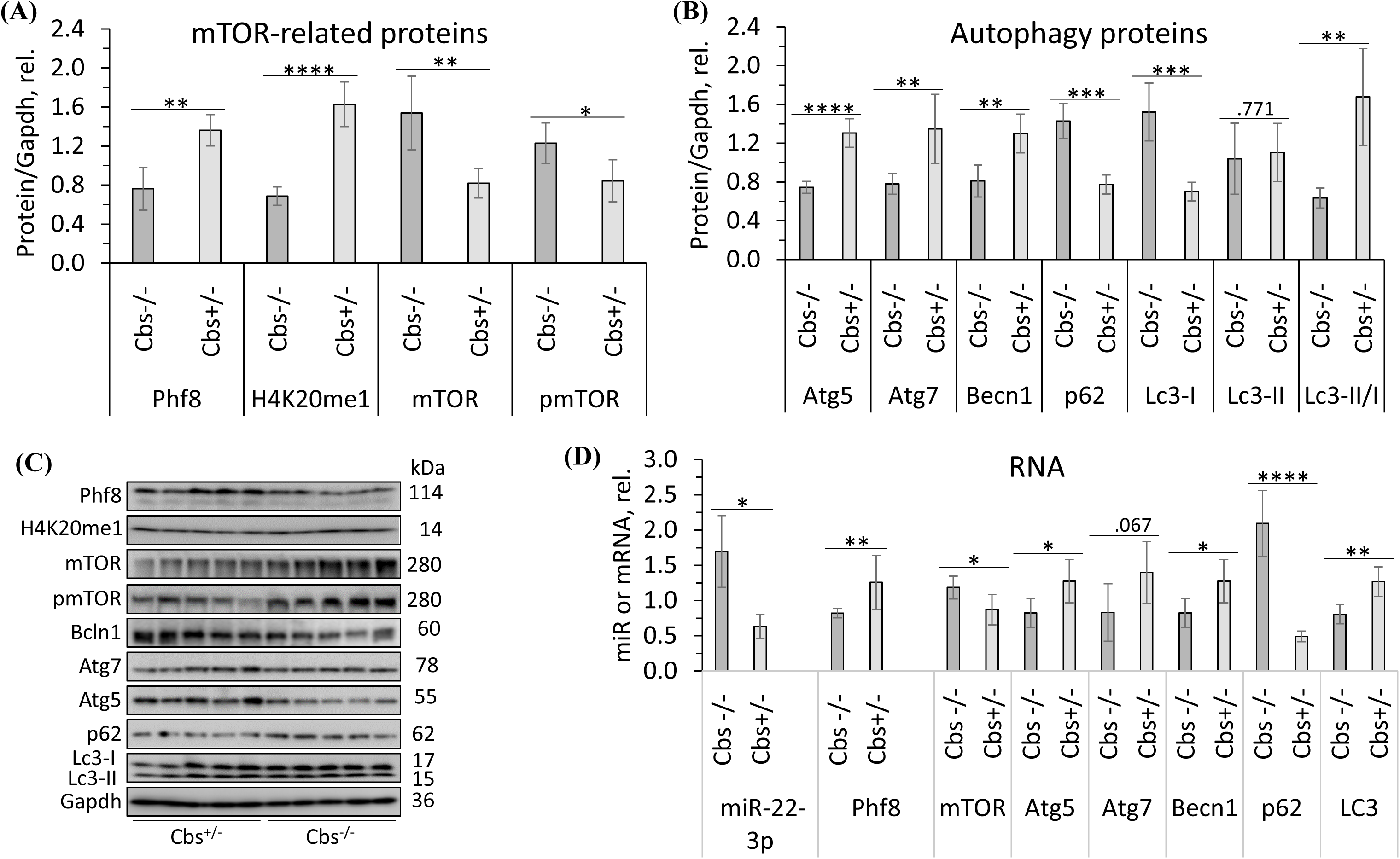
Cbs deficiency affects the expression of miR-22-3p, histone demethylase Phf8, histone H4K20me1 epigenetic mark, mTOR signaling, and autophagy in the mouse heart. Twelve-months-old *Tg-I278T Cbs*^-/-^ mice (n = 5) and their *Tg-I278T Cbs*^+/-^ sibling controls (n = 5) were used in experiments. Bar graphs illustrating quantification of (A) heart miR-22-3p and mRNAs for Phf8, mTOR, Atg, 5, Atg7, Becn1, p62, and Lc3; (B) mTOR signaling-related proteins (Phf8, H4K20me1, mTOR, pmTOR; (C) autophagy-related proteins (Atg5, Atg7, Becn1, p62, Lc3-I, and lipidated Lc3-II) and the Lc3-II/Lc3-I ratio; (D) representative images of western blots used for protein quantification. Indicated RNAs and proteins were quantified by RT-qPCR and western blotting, respectively. Gapdh mRNA and protein were used as references for normalization. *p-* Values were calculated by Mann Whitney test. \**P* < 0.05, \*\**P* < 0.01, \*\*\**P* < 0.001, or \*\*\*\**P* < 0.0001. The numerical values above Atg7 mRNA data in panel (A) and Lc3-I protein data in panel (C) represent *P*-values.

To assess whether the influence of miR-22-3p was propagated downstream of Phf8, we quantified mTOR- and autophagy-related proteins/mRNA. We found that the histone H4K20me1 epigenetic mark was significantly elevated in hearts of *Cbs*^-/-^ mice compared to *Cbs*^+/-^ sibling controls (*P* < 0.0001, Figure 5B). Both mTOR (mRNA, *P* < 0.05, Figure 5A; protein *P* < 0.01, Figure 5B), pmTOR (protein, *P* < 0.05, Figure 5B), and autophagy-related p62 (mRNA, *P* < 0.0001, Figure 5A; protein, *P* < 0.001, Figure 5C) were upregulated while other autophagy-related mRNAs (Atg5, *P* < 0.05; Atg7, *P* < 0.067; Becn1, *P* < 0.05; LC3, *P* < 0.01; Figure 5A) and proteins (Atg5, *P* < 0.0001; Atg7, *P* < 0.001; Becn1, *P* < 0.01; Lc3-I, *P* < 0.001; Figure 5C) were significantly downregulated. Lc3-II protein level was not changed and the Lc3-II/Lc3-I ratio significantly decreased in *Cbs*^-/-^ hearts compared to *Cbs*^+/-^ hearts (*P* < 0.01, Figure 5C). These findings show miR-22-3p mediates the influence of HHcy on Phf8/H4K20me3/mTOR/ autophagy pathway in *Cbs*^-/-^ hearts.

## 4 DISCUSSION

HHcy has been associated with CVD in many studies (38). Endothelial dysfunction plays a central role in CVD and is a common finding in HHcy in humans and in animal models (3-5). To understand the mechanisms by which HHcy disrupts normal cellular function and ultimately causes disease, we used HUVEC, an often-used model of vascular cells (11, 39), to study how Hcy and its metabolites Hcy-thiolactone and *N*-Hcy-protein, all of which accumulate in HHcy, affect the downstream consequences of these effects on mTOR signaling and autophagy, processes important for vascular homeostasis (39).

Our data show that Hcy and its metabolites, Hcy-thiolactone and *N*-Hcy-protein, downregulated the expression of PHF8 by upregulating miR-22-3p and miR-1229-3p. These miRs have not been studied before in the context of Hcy metabolism and it was unknown whether they or PHF8 have any role in vascular endothelial cell function or CVD. We’ve chosen miR-22-3p and miR-1229-3p for investigation because these were the only miRs described in the literature (ref. 24 and 25) as binding to PHF8 3’UTR and on our previous findings that Hcy metabolites inhibit Phf8 in mouse neural cells and brains (ref. 31 and 32).

The downregulation of PHF8 led to elevated levels of H4K20me1, a positive regulator of mTOR expression. The resulting upregulation of mTOR led to the inhibition of autophagy. The ability of Hcy, Hcy-thiolactone, and *N*-Hcy-protein to upregulate miR-22-3p and miR-1229-3p expression and thus affect the PHF8/H4K20me1/mTOR/autophagy pathway, important for vascular homeostasis, can explain the susceptibility of human vascular endothelial cells to HHcy-induced endothelial dysfunction, a harbinger of atherosclerosis.

Studies of metabolic conversions of Hcy that led to the discoveries of Hcy-thiolactone and *N*-Hcy-protein in cultured human fibroblasts, supplied basis for the proposal that these metabolites are responsible for the pathological consequences of HHcy (40). These discoveries were confirmed in HUVEC (11), in mice (34, 41), and in humans (41-43). A recent clinical study has shown that Hcy-thiolactone is a risk predictor of myocardial infarction in patients with coronary artery disease, thereby supplying a support to the hypothesis that Hcy-thiolactone is mechanistically involved in CVD (44, 45). Our present findings that Hcy-thiolactone and *N*-Hcy-protein, as well as Hcy, upregulate miR-22-3p and miR-1229-3p in HUVEC thereby starting a pathway leading to impaired autophagy, suggest that these miRs can also be associated with endothelial dysfunction and supply a new mechanistic explanation of the involvement of HHcy in CVD. Methionine did not affect miR-22-3p and miR-1229-3p, showing that upregulation was specific to these Hcy metabolites.

The effects of Hcy metabolites on gene expression in HUVEC were more pronounced at higher concentrations as shown for H4K20me1, mTOR, BECN1, ATG5 protein (Figure 1 B-G)) and PHF8 mRNA, miR-22-3p, and miR-1229-3p (Figure 2A-C). These findings are in line with the dose effects of HHcy in CVD (46).

HHcy can lead to endothelial dysfunction by other mechanisms such as oxidative stress, production of nitric oxide, oxidized LDL, and inflammation (46). These mechanisms may also be mediated by effects of HHcy on miR-22-3p and miR-1229-3p, which remains to be investigated. Although we have shown that *N*-Hcy-protein, Hcy-thiolactone, and Hcy in HUVEC and genetic HHcy in *Cbs*^-/-^ mice dysregulate PHF8 expression by affecting miR expression in HUVEC it is not excluded that other Hcy metabolites such as S-adenosylhomocysteine (SAH) may also affect miR expression. This possibility remains to be investigated in future studies.

PHF8 is a transcription activator that can bind to promoters of about one-third of human genes (45). Thus, dysregulation of PHF8 expression can affect many biological processes and is likely to cause disease. Indeed, PHF8 depletion has been linked to neuro-logical disorders such as autism spectrum disorder, attention deficit hyperactivity disorder, severe intellectual disability(21, 22) and Alzheimer’s disease (26) while PHF8 up-regulation is a key factor in variety of cancers (24). Our present findings in HUVEC that Hcy metabolites downregulate PHF8 expression via miR-22-3p and miR-1229-3p, resulting in dysregulated mTOR signaling and impaired autophagy flux, suggest that PHF8 depletion can also lead to vascular disease. This suggestion is supported by our present findings that miR-22-3p is upregulated in hearts of Cbs-deficient mice that show impaired mTOR signaling and autophagy flux (Figure 5) in addition to severe HHcy, and endothelial dysfunction (3-5). This suggestion is also supported by findings of other investigators showing that autophagy flux controls endothelial cell homeostasis while impaired autophagy can promote pro-atherogenic phenotype (39).

Each of the three examined metabolites‒*N*-Hcy-protein, Hcy-thiolactone, and Hcy‒ affected the expression of PHF8 mRNA and PHF8-targeting miRs, miR-22-3p and miR-1229-3p (Figure 2A-C). However, effects of *N*-Hcy-protein, Hcy-thiolactone, and Hcy on PHF8 protein and its downstream targets H4K20me1, mTOR, pmTOR (Figure 1A-D), and autophagy related proteins (Figure 1E-K) were metabolite-specific. For example, *N*-Hcy-protein and Hcy-thiolactone significantly downregulated PHF8 and upregulated mTOR and pmTOR (Figure 1A, B, and C, respectively). All three metabolites significantly upregulated H4K20me1 (Figure 1B), and downregulated autophagy related BECN1 (Figure 1E). In contrast, p62 (Figure 1H) and LC3-I (Figure 1J) were not significantly affected by any of these metabolites while other autophagy-related proteins were affected in a metabolite-specific manner. Specifically, ATG7 was downregulated by *N*-Hcy-protein and Hcy-thiolactone but not by Hcy (Figure 1G), ATG5 by N-Hcy-protein and Hcy (Figure 1F), and the autophagy flux LC3-II/LC3-I by Hcy-thiolactone and Hcy but not by *N*-Hcy-protein. These findings suggest that Hcy, Hcy-thiolactone, and *N*-Hcy-protein may influence the expression of mTOR- and autophagy-related proteins independently of their effects on PHF8.

During the past two decades, miRs have been identified as key regulators of the pathophysiological processes involved in atherosclerosis such as signaling (NF-κB, MAPK, SOCS, SDF-1/CXCR4, TGF-β, BMP, TLR) and lipid homeostasis pathways (47). The HDL-carried miRNAs can mediate extracellular signaling by repressing genes in target tissues and HDL interaction with macrophages and endothelial cells may also result in miRNA exchange. Our present findings show that, HHcy-related metabolites, which are associated with CVD, upregulate two miRs targeting PHF8 in HUVEC (Figure 2): miR-22-3p (24) and miR-1229-3p [25] (https://mirtarbase.cuhk.edu.cn/~miRTarBase/miRTarBase_2022/php/detail.php?mirtid=MIRT 036321) (Supplementary Figure S1). Of these, miR-22-3p has been associated with cancer, neuropathy (24), and heart failure (48), while miR-1229-3p has been associated with bladder neoplasms and risk of Alzheimer’s disease (49). We showed that Hcy metabolite-induced miR-22-3p and miR-1229-3p expression lowered histone demethylase PHF8 and upregulated methylated histone H4K20me1, a key regulator that increased mTOR expression, which in turn inhibited autophagy in HUVEC. Notably, we also showed that miR-22-3p was upregulated and that mTOR signaling and autophagy flux were impaired in hearts of Cbs-deficient mice (Figure 5). Taken together, these findings suggest that miR-dependent, Phf8-mediated dysregulation of mTOR signaling pathway and impaired autophagy can be involved in endothelial dysfunction and atherosclerosis induced by HHcy.

The strengths of our study are that it used human vascular endothelial cells and a mouse model in well-designed experiments to address an important area of research relevant to our understanding of endothelial dysfunction and CVD. We elucidated detailed molecular mechanisms and pathways involved, linking Hcy metabolites, miRNAs, PHF8, mTOR, and autophagy. This comprehensive analysis contributes to our understanding of the complex mechanisms underlying endothelial dysfunction and atherosclerosis. The use of miR inhibitors to demonstrate the specific effects of miR-22-3p and miR-1229-3p on PHF8 and downstream signaling, a crucial aspect of the study design, also strengthened the study. The study has also some limitations. For example, we used a single cell line. This, however, is remedied to some extent using a Cbs-deficient mouse model, which allowed us to confirm in vivo our findings in HUVEC. Another limitation is a lack of mechanistic details on how Hcy metabolites induce the expression of miR-22-3p and miR-1229-3p, and how exactly miRs target PHF8 3’UTR, which remains to be addressed in future studies. It would be beneficial to identify and investigate other miRs that target PHF8. The lack of human data is also a limitation that remains to be remedied in future clinical studies. This is important because targeting miR-22-3p and miR-1229-3p (and other miRs) might offer a potentially beneficial therapeutic strategy in CVD.

In conclusion, our findings define a new miR-mediated mechanism involving PHF8 by which HHcy can induce endothelial dysfunction in humans and mice. In this mechanism, *N*-Hcy-protein, Hcy-thiolactone, and Hcy, metabolites associated with HHcy, upregulate the expression of PHF8-targeing miR-22-3p (and miR-1229-3p in humans), which leads to downregulation of PHF8 and dysregulation of PHF8-regulated processes such as mTOR signaling and autophagy.

## ACKNOWLEDGMENTS

Supported in part by grants 2016/23/ N/NZ3/01216, 2018/29/B/NZ4/00771, 2019/33/B/NZ4/01760, 2021/43/B/NZ4/00339 from the National Science Center, and the American Heart Association grant 17GRNT32910002.

## AUTHOR CONTRIBUTION

Ł. Witucki planned, performed and analyzed the biochemical experiments, acquired funding; H. Jakubowski conceived the idea for the project, maintained mice, planned and analyzed experiments, acquired funding, administered the project, and wrote the paper.

## DISCLOSURES

No conflicts of interest, financial or otherwise, are declared by the authors.

## DATA AVAILABILITY STATEMENT

The data that support the findings of this study are available in the methods and/or supplementary material of this article.

## Graphical abstract

Hypothetical pathway leading to autophagy inhibition and endothelial dysfunction in hyperhomocysteinemia. Hcy metabolites upregulate miR22-3p and miR-1229-3p in human endothelial cells and miR22-3p in *Cbs*^-/-^ mouse heart. These miRs downregulate the histone demethylase PHF8, thereby elevating H4K20me1, a positive regulator of mTOR. Upregulation of mTOR and pmTOR inhibits autophagy in human endothelial cells and mouse heart. Up and down arrows show the direction of changes. Hcy, homocysteine; mTOR, mammalian target of rapamycin; pmTOR, phospho-mTOR; PHF8, Plant Homeodomain Finger protein 8.

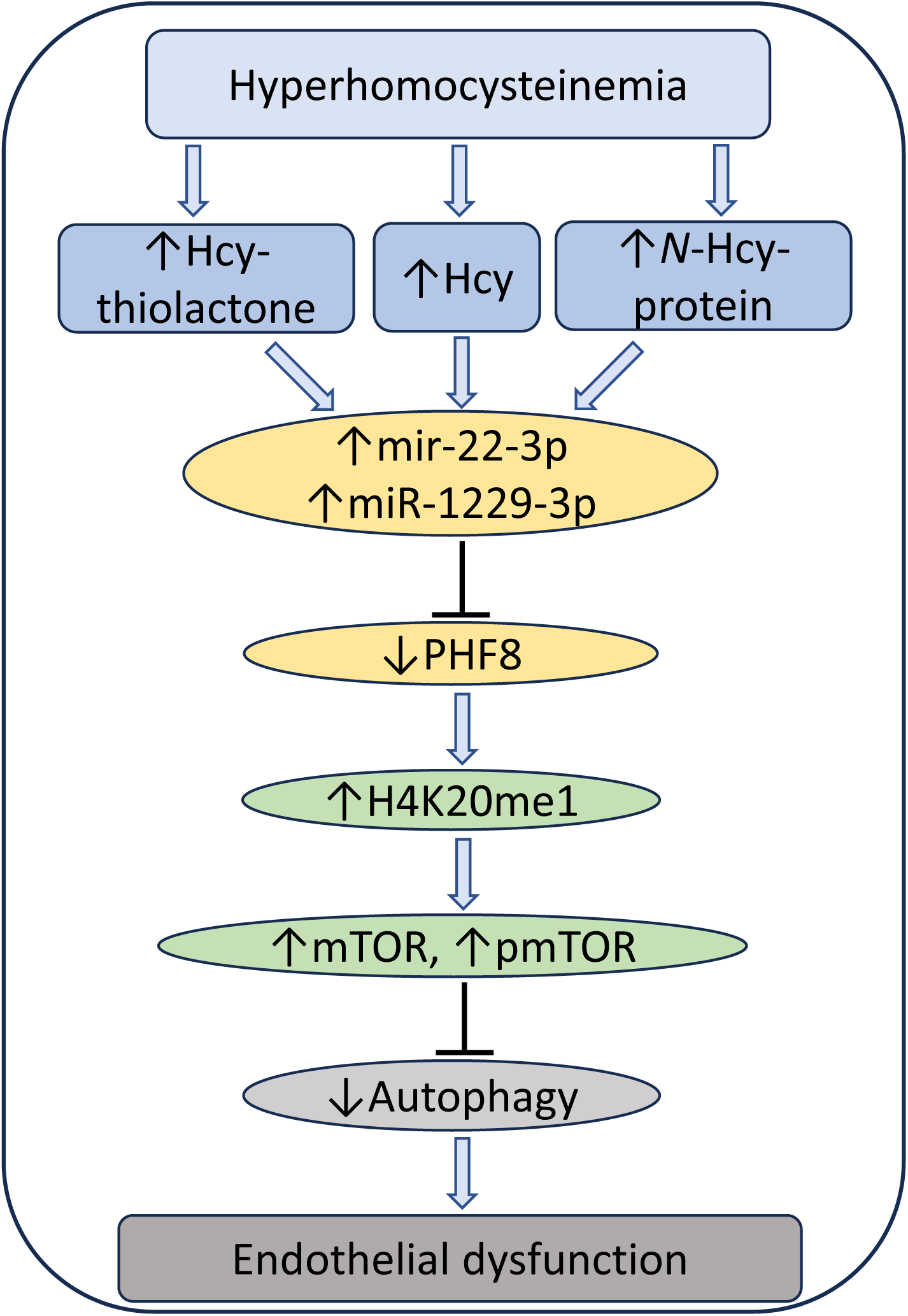

## Notes

### Competing Interest Statement

The authors have declared no competing interest.

### Summary of Updates

The manuscript has been revised to show that our findings are not limited to in vitro cultured human endothelial cells (HUVEC) but are also relevant in vivo in an animal model. This was achieved by including new in vivo data obtained with Cbs-/- that show severe HHcy and endothelial dysfunction, which validated the findings observed in vitro. We have also incuded new data from the dual liciferase reporter experiments to validate miR-22-3p and mir-1229-3p target sites on the PHF8 3'UTR in HUVEC. A graphical abstract has now been included.

## References

1. Libby, P. (2002) Inflammation in atherosclerosis. Nature 420, 868–874

2. Ross, R. (1999) Atherosclerosis--an inflammatory disease. N Engl J Med 340, 115–126

3. Lentz, S. R. (2005) Mechanisms of homocysteine-induced atherothrombosis. J Thromb Haemost 3, 1646–1654

4. Dayal, S., and Lentz, S. R. (2008) Murine models of hyperhomocysteinemia and their vascular phenotypes. Arterioscler Thromb Vasc Biol 28, 1596–1605

5. Esse, R., Barroso, M., Tavares de Almeida, I., and Castro, R. (2019) The Contribution of Homocysteine Metabolism Disruption to Endothelial Dysfunction: State-of-the-Art. Int J Mol Sci 20

6. Poddar, R., Sivasubramanian, N., DiBello, P. M., Robinson, K., and Jacobsen, D. W. (2001) Homocysteine induces expression and secretion of monocyte chemoattractant protein-1 and interleukin-8 in human aortic endothelial cells: implications for vascular disease. Circulation 103, 2717–2723

7. Kerkeni, M., Tnani, M., Chuniaud, L., Miled, A., Maaroufi, K., and Trivin, F. (2006) Comparative study on in vitro effects of homocysteine thiolactone and homocysteine on HUVEC cells: evidence for a stronger proapoptotic and proinflammative homocysteine thiolactone. Mol Cell Biochem 291, 119–126

8. Jakubowski, H. (2019) Homocysteine Modification in Protein Structure/Function and Human Disease. Physiol Rev 99, 555–604

9. Sikora, M., Lewandowska, I., Marczak, L., Bretes, E., and Jakubowski, H. (2020) Cystathionine beta-synthase deficiency: different changes in proteomes of thrombosis-resistant Cbs(-/-) mice and thrombosis-prone CBS(-/-) humans. Sci Rep 10, 10726

10. Sikora, M., and Jakubowski, H. (2021) Changes in redox plasma proteome of Pon1-/- mice are exacerbated by a hyperhomocysteinemic diet. Free Radic Biol Med 169, 169–180

11. Jakubowski, H., Zhang, L., Bardeguez, A., and Aviv, A. (2000) Homocysteine thiolactone and protein homocysteinylation in human endothelial cells: implications for atherosclerosis. Circ Res 87, 45–51

12. Gurda, D., Handschuh, L., Kotkowiak, W., and Jakubowski, H. (2015) Homocysteine thiolactone and N-homocysteinylated protein induce pro-atherogenic changes in gene expression in human vascular endothelial cells. Amino Acids 47, 1319–1339

13. Olejniczak, M., Urbanek, M. O., Jaworska, E., Witucki, L., Szczesniak, M. W., Makalowska, I., and Krzyzosiak, W. J. (2016) Sequence-non-specific effects generated by various types of RNA interference triggers. Biochim Biophys Acta 1859, 306–314

14. Starega-Roslan, J., and Krzyzosiak, W. J. (2013) Analysis of microRNA length variety generated by recombinant human Dicer. Methods Mol Biol 936, 21–34

15. Starega-Roslan, J., Galka-Marciniak, P., and Krzyzosiak, W. J. (2015) Nucleotide sequence of miRNA precursor contributes to cleavage site selection by Dicer. Nucleic Acids Res 43, 10939-10951

16. Starega-Roslan, J., Koscianska, E., Kozlowski, P., and Krzyzosiak, W. J. (2011) The role of the precursor structure in the biogenesis of microRNA. Cell Mol Life Sci 68, 2859–2871

17. Stroynowska-Czerwinska, A., Fiszer, A., and Krzyzosiak, W. J. (2014) The panorama of miRNA-mediated mechanisms in mammalian cells. Cell Mol Life Sci 71, 2253–2270

18. Minjares, M., Wu, W., and Wang, J. M. (2023) Oxidative Stress and MicroRNAs in Endothelial Cells under Metabolic Disorders. Cells 12

19. Zhou, S. S., Jin, J. P., Wang, J. Q., Zhang, Z. G., Freedman, J. H., Zheng, Y., and Cai, L. (2018) miRNAS in cardiovascular diseases: potential biomarkers, therapeutic targets and challenges. Acta Pharmacol Sin 39, 1073–1084

20. Mens, M. M. J., Heshmatollah, A., Fani, L., Ikram, M. A., Ikram, M. K., and Ghanbari, M. (2021) Circulatory MicroRNAs as Potential Biomarkers for Stroke Risk: The Rotterdam Study. Stroke 52, 945–953

21. Sobering, A. K., Bryant, L. M., Li, D., McGaughran, J., Maystadt, I., Moortgat, S., Graham, J. M., Jr., van Haeringen, A., Ruivenkamp, C., Cuperus, R., Vogt, J., Morton, J., Brasch-Andersen, C., Steenhof, M., Hansen, L. K., Adler, E., Lyonnet, S., Pingault, V., Sandrine, M., Ziegler, A., Donald, T., Nelson, B., Holt, B., Petryna, O., Firth, H., McWalter, K., Zyskind, J., Telegrafi, A., Juusola, J., Person, R., Bamshad, M. J., Earl, D., University of Washington Center for Mendelian, G., Tsai, A. C., Yearwood, K. R., Marco, E., Nowak, C., Douglas, J., Hakonarson, H., and Bhoj, E. J. (2022) Variants in PHF8 cause a spectrum of X-linked neurodevelopmental disorders and facial dysmorphology. HGG Adv 3, 100102

22. Laumonnier, F., Holbert, S., Ronce, N., Faravelli, F., Lenzner, S., Schwartz, C. E., Lespinasse, J., Van Esch, H., Lacombe, D., Goizet, C., Phan-Dinh Tuy, F., van Bokhoven, H., Fryns, J. P., Chelly, J., Ropers, H. H., Moraine, C., Hamel, B. C., and Briault, S. (2005) Mutations in PHF8 are associated with X linked mental retardation and cleft lip/cleft palate. J Med Genet 42, 780–786

23. Chen, X., Wang, S., Zhou, Y., Han, Y., Li, S., Xu, Q., Xu, L., Zhu, Z., Deng, Y., Yu, L., Song, L., Chen, A. P., Song, J., Takahashi, E., He, G., He, L., Li, W., and Chen, C. D. (2018) Phf8 histone demethylase deficiency causes cognitive impairments through the mTOR pathway. Nat Commun 9, 114

24. Cai, M. Z., Wen, S. Y., Wang, X. J., Liu, Y., and Liang, H. (2020) MYC Regulates PHF8, Which Promotes the Progression of Gastric Cancer by Suppressing miR-22-3p. Technol Cancer Res Treat 19, 1533033820967472

25. Helwak, A., Kudla, G., Dudnakova, T., and Tollervey, D. (2013) Mapping the human miRNA interactome by CLASH reveals frequent noncanonical binding. Cell 153, 654–665

26. Witucki, L., and Jakubowski, H. (2023) Depletion of Paraoxonase 1 (Pon1) Dysregulates mTOR, Autophagy, and Accelerates Amyloid Beta Accumulation in Mice. Cells 12

27. Kaldirim, M., Lang, A., Pfeiler, S., Fiegenbaum, P., Kelm, M., Bonner, F., and Gerdes, N. (2022) Modulation of mTOR Signaling in Cardiovascular Disease to Target Acute and Chronic Inflammation. Front Cardiovasc Med 9, 907348

28. Samidurai, A., Kukreja, R. C., and Das, A. (2018) Emerging Role of mTOR Signaling-Related miRNAs in Cardiovascular Diseases. Oxid Med Cell Longev 2018, 6141902

29. Mei, X., Qi, D., Zhang, T., Zhao, Y., Jin, L., Hou, J., Wang, J., Lin, Y., Xue, Y., Zhu, P., Liu, Z., Huang, L., Nie, J., Si, W., Ma, J., Ye, J., Finnell, R. H., Saiyin, H., Wang, H., Zhao, J., Zhao, S., and Xu, W. (2020) Inhibiting MARSs reduces hyperhomocysteinemia-associated neural tube and congenital heart defects. EMBO Mol Med 12, e9469

30. Khayati, K., Antikainen, H., Bonder, E. M., Weber, G. F., Kruger, W. D., Jakubowski, H., and Dobrowolski, R. (2017) The amino acid metabolite homocysteine activates mTORC1 to inhibit autophagy and form abnormal proteins in human neurons and mice. FASEB J 31, 598–609

31. Witucki, L., and Jakubowski, H. (2023) Homocysteine metabolites inhibit autophagy, elevate amyloid beta, and induce neuropathy by impairing Phf8/H4K20me1-dependent epigenetic regulation of mTOR in cystathionine beta-synthase-deficient mice. J Inherit Metab Dis

32. Witucki, L., Borowczyk, K., Suszynska-Zajczyk, J., Warzych, E., Pawlak, P., and Jakubowski, H. (2023) Deletion of the Homocysteine Thiolactone Detoxifying Enzyme Bleomycin Hydrolase, in Mice, Causes Memory and Neurological Deficits and Worsens Alzheimer’s Disease-Related Behavioral and Biochemical Traits in the 5xFAD Model of Alzheimer’s Disease. J Alzheimers Dis

33. Gupta, S., Kuhnisch, J., Mustafa, A., Lhotak, S., Schlachterman, A., Slifker, M. J., Klein-Szanto, A., High, K. A., Austin, R. C., and Kruger, W. D. (2009) Mouse models of cystathionine beta-synthase deficiency reveal significant threshold effects of hyperhomocysteinemia. Faseb J 23, 883–893

34. Jakubowski, H., Perla-Kajan, J., Finnell, R. H., Cabrera, R. M., Wang, H., Gupta, S., Kruger, W. D., Kraus, J. P., and Shih, D. M. (2009) Genetic or nutritional disorders in homocysteine or folate metabolism increase protein N-homocysteinylation in mice. Faseb J 23, 1721–1727

35. Jakubowski, H. (2020) Proteomic exploration of cystathionine beta-synthase deficiency: implications for the clinic. Expert Rev Proteomics 17, 751–765

36. Witucki, L., Kurpik, M., Jakubowski, H., Szulc, M., Lukasz Mikolajczak, P., Jodynis-Liebert, J., and Kujawska, M. (2022) Neuroprotective Effects of Cranberry Juice Treatment in a Rat Model of Parkinson’s Disease. Nutrients 14

37. Livak, K. J., and Schmittgen, T. D. (2001) Analysis of relative gene expression data using real-time quantitative PCR and the 2(-Delta Delta C(T)) Method. Methods 25, 402–408

38. Smith, A. D., and Refsum, H. (2021) Homocysteine - from disease biomarker to disease prevention. J Intern Med 290, 826–854

39. Mameli, E., Martello, A., and Caporali, A. (2022) Autophagy at the interface of endothelial cell homeostasis and vascular disease. FEBS J 289, 2976–2991

40. Jakubowski, H. (1997) Metabolism of homocysteine thiolactone in human cell cultures. Possible mechanism for pathological consequences of elevated homocysteine levels. J Biol Chem 272, 1935–1942

41. Chwatko, G., Boers, G. H., Strauss, K. A., Shih, D. M., and Jakubowski, H. (2007) Mutations in methylenetetrahydrofolate reductase or cystathionine beta-synthase gene, or a high-methionine diet, increase homocysteine thiolactone levels in humans and mice. Faseb J 21, 1707–1713

42. Bogdanski, P., Pupek-Musialik, D., Dytfeld, J., Lacinski, M., Jablecka, A., and Jakubowski, H. (2008) Plasma homocysteine is a determinant of tissue necrosis factor-alpha in hypertensive patients. Biomed Pharmacother 62, 360–365

43. Perla-Kajan, J., Stanger, O., Luczak, M., Ziolkowska, A., Malendowicz, L. K., Twardowski, T., Lhotak, S., Austin, R. C., and Jakubowski, H. (2008) Immunohistochemical detection of N-homocysteinylated proteins in humans and mice. Biomed Pharmacother 62, 473–479

44. Borowczyk, K., Piechocka, J., Glowacki, R., Dhar, I., Midtun, O., Tell, G. S., Ueland, P. M., Nygard, O., and Jakubowski, H. (2019) Urinary excretion of homocysteine thiolactone and the risk of acute myocardial infarction in coronary artery disease patients: the WENBIT trial. J Intern Med 285, 232–244

45. Qi, H. H., Sarkissian, M., Hu, G. Q., Wang, Z., Bhattacharjee, A., Gordon, D. B., Gonzales, M., Lan, F., Ongusaha, P. P., Huarte, M., Yaghi, N. K., Lim, H., Garcia, B. A., Brizuela, L., Zhao, K., Roberts, T. M., and Shi, Y. (2010) Histone H4K20/H3K9 demethylase PHF8 regulates zebrafish brain and craniofacial development. Nature 466, 503–507

46. Gueant, J. L., Gueant-Rodriguez, R. M., Oussalah, A., Zuily, S., and Rosenberg, I. (2023) Hyperhomocysteinemia in Cardiovascular Diseases: Revisiting Observational Studies and Clinical Trials. Thromb Haemost 123, 270–282

47. Feinberg, M. W., and Moore, K. J. (2016) MicroRNA Regulation of Atherosclerosis. Circ Res 118, 703–720

48. Lu, W., Liu, X., Zhao, L., Yan, S., Song, Q., Zou, C., and Li, X. (2023) MiR-22-3p in exosomes increases the risk of heart failure after down-regulation of FURIN. Chem Biol Drug Des 101, 550-567

49. Ghanbari, M., Ikram, M. A., de Looper, H. W. J., Hofman, A., Erkeland, S. J., Franco, O. H., and Dehghan, A. (2016) Genome-wide identification of microRNA-related variants associated with risk of Alzheimer’s disease. Sci Rep 6, 28387

