## Supplementary for "Homocysteine Metabolites Impair the PHF8/H4K20me1/mTOR/Autophagy Pathway by Upregulating the Expression of Histone Demethylase PHF8-targeting microRNAs in Human Vascular Endothelial Cells and Mice"

**Supplementary Material**

**Supplementary Figure S1**

**Supplementary Figure S2**

**Supplementary Table S1**

**Supplementary Table S2**

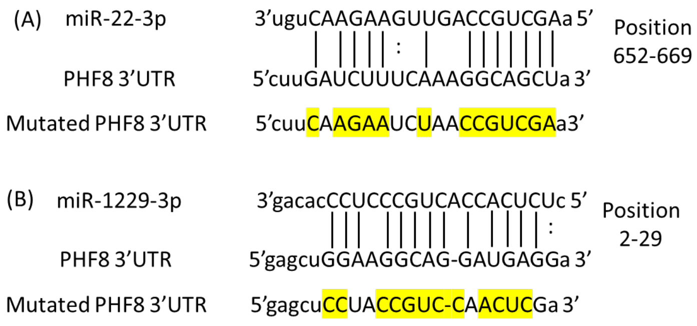


**Supplementary Figure S1.** Validated PHF8 3′UTR target sites for miR-22-3p (A) and miR-1229-3p (B). The PHF8 site for miR-22-3p (position 652 – 669) has been validated by the dual-luciferase reporter vector assay (ref. 24 and this work). The PHF8 site for miR-1229-3p (position 2 – 23) has been validated by miR-mRNA specific interaction using the crosslinking, ligation, and sequencing of hybrids (CLASH) method (ref. 25) and dual-luciferase reporter vector assay (this work). Mutated nucleotides in PHF8 3′UTR are shown by yellow highlighting.

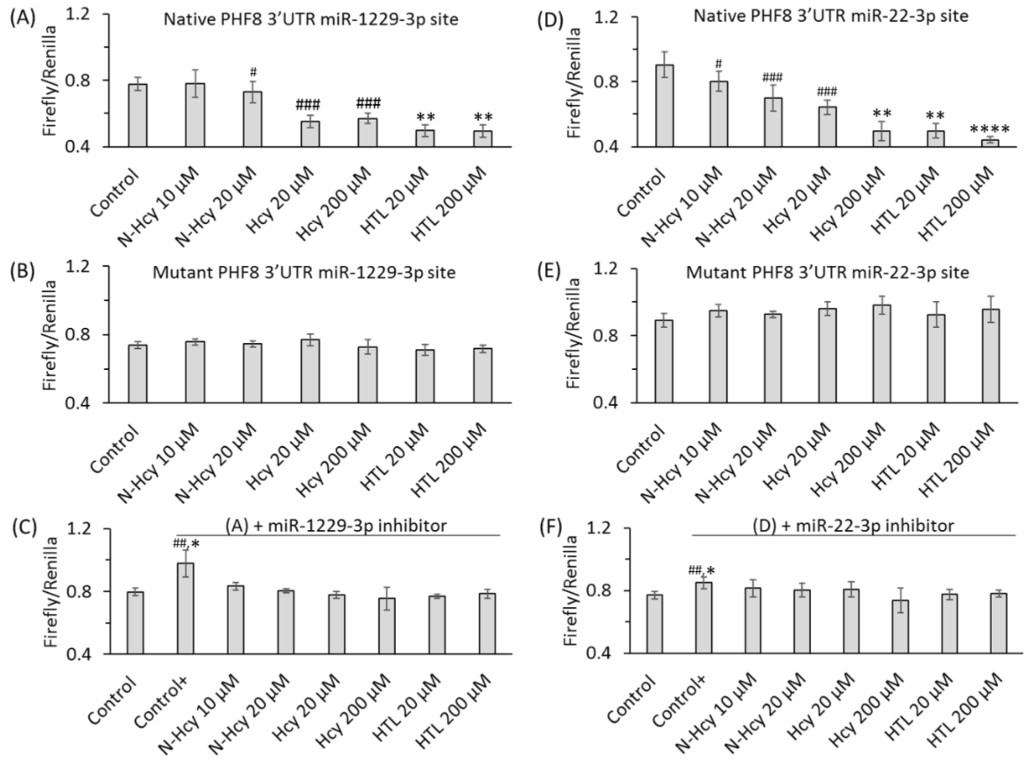


**Supplementary Figure S2.** Binding of miR-1229-3p and miR-22-3p to PHF8 3'UTR in HUVEC confirmed by the dual luciferase assay. The 3' UTRs fragments of the human *PHF8* gene having native or mutated miR-22-3p or miR-1229-3p binding sites were cloned into the pmirGLO vector (Promega) cut with XbaI and DraI restriction enzymes (New England BioLabs). HUVEC cells were seeded on 96-well plate (15,000 cells/well), grown to 70% confluency (~24 h), and transfected with a *PHF8 3'UTR*-containg plasmid (0.01 µg) in the absence and presence of a specific miR inhibitor (0.5 nM) for 4 h using Lipofectamine 2000 (Invitrogen). HUVEC monolayers were rinsed twice with PBS, overlaid with M199 medium without methionine (Thermo Scientific) containing 5% dialyzed FBS (Millipore Sigma) and untreated (Control, Control+) or treated with *N*-Hcy-protein, Hcy-thiolactone, or Hcy for 24 h. The firefly and renilla luminescence were quantified using a Dual-Glo® Luciferase Assay System (Promega, USA) and the firefly/renilla luminescence ratios calculated. HUVEC cells transfected with (A) PHF8 3'UTR containing native binding site for miR-1229-3p, (B) PHF8 3'UTR containing mutated binding site for miR-1229-3p, (C) miR-1229-3p inhibitor and PHF8 3'UTR containing native binding site for miR-1229-3p, (D) PHF8 3'UTR containing native binding site for miR-22-3p, (E) PHF8 3'UTR containing mutated binding site for miR-22-3p, (F) miR-22-3p inhibitor and PHF8 3'UTR containing native binding site for miR-22-3p. T test: $^{\#\#}P < 0.01$, $^{\#\#\#}P < 0.001$. Mann-Whitney test: $^{*}P<0.05$, $^{**}P<0.01$, or $^{****}P<0.0001$.

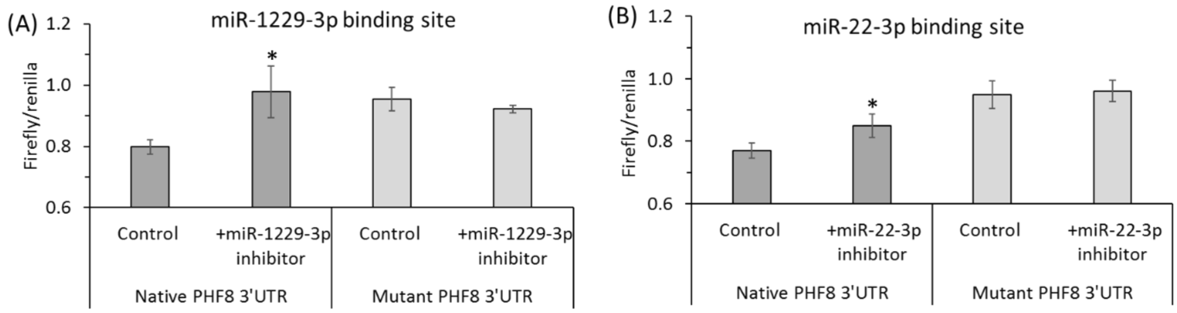


**Supplementary Figure S3.** Inhibitors of miR-1229-3p or miR-22-3p stimulate the activity of native but not mutant PHF8 3'UTR in a dual luciferase assay. HUVEC were transfected with PHF8 3'UTR containing native or mutated binding site for miR-1229-3p or miR-22-3p in the absence (A) or presence (B) of miR-1229-3p inhibitor or miR-22-3p inhibitor, respectively. *P*-values were calculated by Mann Whitney test. *$p<0.05$.

| **Supplementary Table S1** Primers used for RT-qPCR | |
|---|---|
| ***Gene*** | Primer sequence |
| *PHF8* | Forward: 5′-TGGGAGCATGCTTCAAGG-3′ |
| | Reverse: 5′-GATTTCAAAGCAGGGTCATCA-3′ |
| *GAPDH* | Forward: 5′-AGCCACATCGCTCAGACAC-3′ |
| | Reverse: 5′-GCCAATACGACCAAATCC-3′ |
| *mTOR* | Forward: 5′-GCAGCTGCATGGGGTTTA-3′ |
| | Reverse: 5′-CCCGAGGGATCATACAGGT-3′ |
| *ATG5* | Forward: 5′-AGTAGTTGCCTGGAGGAGCG-3′ |
| | Reverse: 5′-GTCTCGCCAACTCCACCTTG-3′ |
| *ATG7* | Forward: 5′-GATGAAGCTCCCAAGGACAT-3′ |
| | Reverse: 5′-GTGGGAGCACTCATGTCAAA-3′ |
| *BECN1* | Forward: 5′-GGGCTCCCGAGGGATGG-3′ |
| | Reverse: 5′-TTCCTCCTGGGTCTCTCCTG-3′ |
| *LC3* | Forward: 5′-CATGAGCGAGTTGGTCAAGA-3′ |
| | Reverse: 5′-CCATGCTGTGCTGGTTCA-3′ |
| *P62* | Forward: 5′-GGTCGCGCTCACCTTTCT-3′ |
| | Reverse: 5′-TCCTTTCTCAAGCCCCATGTT-3′ |
| *miR-22-3p* | 5′-AAGCTGCCAGTTGAAGAACTGT-3′ |
| *miR-1229-3p* | 5′-CTCTCACCACTGCCCTCCCACAG-3′ |
| *18s rRNA* | Reverse: 5′-AGGAATTCCCAGTAAGTGCG-3′ |
| | Reverse: 5′-GCCTCACTAAACCATCCAA-3′ |
| *U6 snRNA* | Reverse: 5′-CTCGCTTCGGCAGCACA-3′ |
| | Reverse: 5′-AACGCTTCACGAATTTGCGT-3′ |

**Supplementary Table S2.** The inhibitor of miR-22-3p abrogates effects of Hcy metabolites on the expression of PHF8, H4K20me1, mTOR-, and autophagy-related proteins/mRNAs in HUVEC *.

| Metabolite | | PHF8 | | | H4K20me1 | | | mTOR | | | pmTOR | | |
|---|---|---|---|---|---|---|---|---|---|---|---|---|---|
| | | -inhibitor | +inhibitor | *P*-value | -inhibitor | +inhibitor | *P*-value | -inhibitor | +inhibitor | *P*-value | -inhibitor | +inhibitor | *P*-value |
| None | protein | 1.00±0.11 | 1.27±0.08 | 0.001 | 1.00±0.18 | 0.77±0.13 | 0.002 | 1.00±0.18 | 0.80±0.05 | 0.009 | 1.00±0.32 | 0.80±0.05 | 0.009 |
| | mRNA | 1.00±0.13 | 2.32±0.35 | 0.0001 | | | | 1.00±0.22 | 0.57±0.15 | 0.002 | | | |
| N-Hcy-protein | protein | 0.64±0.05 | 0.98±0.07 | 0.002 | 1.72±0.14 | 0.94±0.05 | 0.001 | 1.64±0.11 | 0.95±0.06 | 0.001 | 1.50±0.08 | 0.95±0.06 | 0.006 |
| | mRNA | 0.46±0.01 | 1.12±0.05 | 3E-05 | | | | 1.62±0.17 | 0.88±0.29 | 0.020 | | | |
| Hcy-thiolactone | protein | 0.71±0.10 | 1.04±0.05 | 0.007 | 1.66±0.12 | 0.94±0.05 | 0.001 | 1.68±0.13 | 0.95±0.07 | 0.001 | 1.40±0.13 | 0.95±0.07 | 0.001 |
| | mRNA | 0.53±0.07 | 1.48±0.29 | 0.006 | | | | 1.74±0.10 | 0.79±0.06 | 0.0001 | | | |
| Hcy | protein | 0.67±0.09 | 1.03±0.05 | 0.004 | 1.51±0.15 | 0.90±0.05 | 0.002 | 1.55±0.25 | 0.85±0.10 | 0.010 | 1.57±0.10 | 1.06±0.05 | 0.001 |
| | mRNA | 0.54±0.06 | 1.38±0.22 | 0.003 | | | | 1.52±0.18 | 0.82±0.11 | 0.005 | | | |
| | | ATG5 | | | ATG7 | | | BECN1 | | | LC3 | | |
| | | -inhibitor | +inhibitor | *P* value | -inhibitor | +inhibitor | *P* value | -inhibitor | +inhibitor | *P* value | -inhibitor | +inhibitor | *P* value |
| None | Protein[#] | 1.00±0.05 | 1.19±0.15 | 0.018 | 1.00±0.08 | 1.26±0.09 | 0.001 | 1.00±0.06 | 1.12±0.09 | 0.021 | 1.00±0.12 | 0.97±0.08 | 0.598 |
| | mRNA | 1.00±0.16 | 1.36±0.20 | 0.006 | 1.00±0.16 | 1.43±0.21 | 0.004 | 1.00±0.17 | 0.57±0.15 | 0.002 | 1.00±0.15 | 1.36±0.35 | 0.043 |
| N-Hcy-protein | protein | 0.77±0.04 | 0.98±0.07 | 0.002 | 0.77±0.04 | 1.10±0.06 | 0.001 | 0.75±0.05 | 1.02±0.07 | 0.006 | 0.69±0.05 | 0.91±0.06 | 0.009 |
| | mRNA | 0.57±0.04 | 1.10±0.12 | 0.002 | 0.63±0.09 | 1.17±0.08 | 0.003 | 0.63±0.09 | 1.00±0.05 | 0.003 | 0.69±0.05 | 1.05±0.10 | 0.006 |
| Hcy-thiolactone | protein | 0.77±0.04 | 0.98±0.07 | 0.002 | 0.77±0.04 | 1.10±0.06 | 0.001 | 0.61±0.06 | 1.05±0.05 | 0.001 | 1.00±0.12 | 1.19±0.24 | 0.026 |
| | mRNA | 0.46±0.01 | 1.12±0.05 | 3E-05 | 0.82±0.13 | 1.25±0.06 | 0.007 | 0.87±0.08 | 1.14±0.17 | 0.066 | 0.79±0.18 | 1.24±0.01 | 0.019 |
| Hcy | protein | 0.73±0.09 | 1.03±0.18 | 0.063 | 0.73±0.19 | 1.07±0.07 | 0.039 | 0.68±0.06 | 1.01±0.06 | 0.003 | 0.69±0.07 | 1.07±0.14 | 0.013 |
| | mRNA | 0.46±0.01 | 1.12±0.05 | 3E-05 | 0.75±0.06 | 1.24±0.10 | 0.002 | 0.86±0.10 | 1.02±0.10 | 0.009 | 0.78±0.06 | 1.27±0.05 | 0.0004 |

* HUVEC were transfected with miR-22-3p inhibitor. Transfected (+ inhibitor) and nontransfected (- inhibitor) HUVEC were incubated with 20 μM of indicated Hcy metabolites. PHF8, H4K20me1, mTOR- mRNA and protein, and pmTOR protein were quantified by RT-qPCR and western blotting, respectively. [#] For LC3 protein, values for LC3-II/LC3-I ratio are shown. The data for (- inhibitor) and (+ inhibitor) are illustrated graphically in Figure 1 and Figure 3, respectively. Two-sided *T*-test *P*-values are shown.
